# Neuroimmune characterization of optineurin insufficiency mouse model during ageing

**DOI:** 10.1101/2023.03.09.531905

**Authors:** Nikolina Mohovic, Josip Peradinovic, Andrea Markovinovic, Raffaello Cimbro, Zeljka Minic, Marin Dominovic, Hrvoje Jakovac, Jerneja Nimac, Boris Rogelj, Ivana Munitic

## Abstract

Optineurin is a multifunctional polyubiquitin-binding protein implicated in inflammatory signalling. Optineurin mutations are associated with amyotrophic lateral sclerosis (ALS) and frontotemporal dementia (FTD), neurodegenerative diseases characterised by neuronal loss, neuroinflammation, and peripheral immune disbalance. However, the pathogenic role of optineurin mutations is unclear. We previously observed no phenotype in the unmanipulated young optineurin insufficiency mice (Optn^470T^), designed to mimic ALS/FTD-linked truncations deficient in polyubiquitin binding. The purpose of this study was to investigate whether ageing would trigger neurodegeneration. We performed a neuroimmune characterization of ageing wild-type (WT) and Optn^470T^ mice. No motor or cognitive differences were detected between the genotypes. Neuropathological analyses demonstrated signs of ageing including lipofuscin accumulation and microglial activation. However, this was not worsened in Optn^470T^ mice, and they did not exhibit TAR DNA-binding protein 43 (TDP-43) aggregation or neuronal loss. Spleen immunophenotyping uncovered T cell immunosenescence at two years but without notable differences between the WT and Optn^470T^ mice. Conventional dendritic cells (cDC) and macrophages exhibited increased expression of activation markers in two-year-old Optn^470T^ males but not females, although the numbers of innate immune cells were similar between genotypes. Altogether, a combination of optineurin insufficiency and ageing did not induce ALS/FTD-like neuropathology in mice.

## Introduction

Amyotrophic lateral sclerosis (ALS) is a fatal neurodegenerative disease that causes motor neuron loss in the brain, brainstem, and spinal cord (1). It is characterized by an exceptionally high clinical and genetic heterogeneity and influenced by still unknown environmental factors. Mutations in > 50 genes have been found in both patients with and without a family history of the disease. The most frequently affected genes are chromosome 9 open reading frame 72 (*C9ORF72*), TAR DNA-binding protein 43 (*TARDBP*), superoxide dismutase 1 (*SOD1*), and fused in sarcoma (*FUS*), which are implicated in the regulation of the DNA/RNA metabolism, nucleocytoplasmic and vesicular transport, and oxidative stress (2,3). Notably, mutations in these genes also directly increase protein aggregation, a key hallmark of all neurodegenerative diseases, with TAR DNA-binding protein 43 (TDP-43) aggregates found in > 95% of ALS patients (4,5). Other pathogenic mechanisms have been proposed including dysregulated inflammatory signalling linked to mutations in genes encoding for TANK-binding kinase 1 (*TBK1*), cylindromatosis (*CYLD*), and optineurin (*OPTN*) (6–9), which could primarily affect the immune system and elicit secondary neuronal damage. A substantial fraction of ALS patients develops cognitive symptoms and frontotemporal dementia (FTD), characterized by neuronal loss in frontal and temporal lobes (10). Notably, mutations in *TBK1*, *CYLD*, and *OPTN* have been identified in both ALS and FTD patients, supporting the idea that these disorders are two different manifestations of a single clinicopathological spectrum (11–13,7,14).

Optineurin, encoded by the *OPTN* gene, is a multifunctional polyubiquitin-binding adaptor protein found in high levels in several organs including the brain, spinal cord, spleen, heart, liver, lung, and skeletal muscles (15). Optineurin specifically binds to methionine 1- and lysine 63-linked polyubiquitin chains via its C-terminal ubiquitin-binding region (16–18). It has been reported to regulate several cellular processes, including inflammatory signalling, autophagy, vesicle trafficking, and cell death, but the precise role of optineurin mutations in ALS and/or FTD pathogenesis is puzzling and has been shown to vary in different experimental settings (20). Several mouse models were generated to investigate the role of optineurin in neurodegeneration. One report detected slight motor deficits (in vertical rearing activity) and dysmyelination in the spinal cords of optineurin deficient (Optn^-/-^) mice at three months of age, with no further phenotype exacerbation up to two years, and no motor neuron loss (15). However, motor deficits were not detected in two other Optn^-/-^ mouse models (21,22), despite TDP-43 aggregation and diminished numbers of spinal cord neurons detected in one of the studies (21). Notably, no signs of neuroinflammation or protein aggregation were observed in one-year-old mice with CNS-specific optineurin knock-out (23). Similarly, models carrying optineurin C-terminal truncation (Optn^470T^) or point mutation (Optn^D477N^), mimicking some ALS patient mutations that disrupt ubiquitin-binding, showed no overt neuroinflammation or ALS pathology (at two months, and one year, respectively) (17,24). Isolated primary microglia from the Optn^470T^ mouse model showed decreased activation of TBK1 and subsequent interferon (IFN)-β production (24), but no defect in phagocytosis (25) or TDP-43 aggregation (26). However, detailed characterisation and/or long-term repercussions of the ubiquitin-binding optineurin mutations in the *in vivo* models are lacking so it is unknown if these models are similar or distinct to Optn^-/-^ mice. This is of particular interest because even though most studies argue for a loss-of-function mechanism of optineurin mutations in ALS/FTD (19), certain differences were reported between carriers of heterozygous E478G mutation and homozygous Q398X truncation (21), suggesting a potential distinct mechanism of individual optineurin mutations.

Ageing is a major risk factor for all neurodegenerative diseases (27). In the immune system, it is marked by a lower capacity to cope with various exogenous and endogenous stressors, which ultimately leads to a higher inflammatory response. The resulting low-grade chronic inflammation has been termed inflammageing (28). This is accompanied by alterations in both innate and adaptive immune cell numbers, including a decrease in naïve and an increase in memory T cells, and functional defects in phagocytosis (29). Altogether these changes, also known as immunosenescence, can be detected in both mice and humans with very few differences. In the CNS, inflammageing leads to higher astrocyte and microglial activation accompanied by decreased phagocytosis (30). Neurons are particularly sensitive to high concentrations of inflammatory mediators, impaired debris clean-up, and/or lack of trophic support by glia, which trigger the so-called non-cell autonomous neuronal death, the most common type of cell death occurring during the neurodegenerative process.

We have previously observed proinflammatory and anti-inflammatory factor disbalance in lipopolysaccharide-stimulated macrophages and microglia from the Optn^470T^ mouse model, which was designed to mimic C-terminal truncations found in ALS/FTD patients (31,24). This opened the possibility that a primary defect in the immune system drives neurodegeneration. However, we found no overt neurological phenotype in the unmanipulated young Optn^470T^ mice. Since ageing is a key risk factor for neurodegeneration, in this study we analysed if a combination of two hits - optineurin mutation and ageing - would accelerate immunosenescence and/or inflammageing to uncover ALS or FTD-like neuropathology. To this end, we compared motor and cognitive neurological symptoms and performed an extensive characterisation of various aspects of neurodegeneration and neuroinflammation, including microgliosis, astrocytosis, TDP-43 aggregation, neuronal loss, and the innate and adaptive immune system activation in WT and Optn^470T^ mice up to two years of age.

## Materials and methods

### Animals

C57BL/6 mice were purchased from Jackson and expanded in the animal facility at the Medical School, University of Rijeka. Optn^470T^ mice were generated as previously described (31) and backcrossed to C57BL/6 genetic background 11 times and subsequently crossed among themselves to obtain homozygous mice used in this study. Six months, one- and two-year-old Optn^470T^ mice were used for experiments. Sex- and age-matched C57BL/6 mice (hereafter called WT) were used as controls. Mice were kept as recommended by the institutional and national guidelines. Experimental procedures were performed according to the Animal Research: Reporting of In Vivo Experiments (ARRIVE) guidelines and the European Communities Council Directive of 24 November 1986 (86/609/EEC). They were approved by the Ethics Committee of the Department of Biotechnology and Medical School of the University of Rijeka, and the Ministry of Agriculture of the Republic of Croatia.

### Reagents

Primary antibodies used for immunofluorescence or western blot analyses were goat anti-GFAP (#ab53554) from Abcam (Cambridge, UK), rabbit anti-Iba1 (#019-19741) from Wako (Osaka, Japan), mouse anti-NeuN (#MAB377) from Merck Millipore (Burlington, MA, USA), rabbit N-terminal anti-TDP-43 (#10782-2-AP) and rabbit C-terminal anti-TDP-43 (#12892-1-AP) from Proteintech (Manchester, UK); goat anti-ChAT (#AB144P), mouse anti-β-tubulin (#T8328) and rabbit anti-phospho-TDP-43 (#SAB4200223) from Sigma Aldrich (St. Louis, MO, USA). Secondary antibodies labelled with horseradish peroxidase (HRP; anti-rabbit #111-035-144 and anti-mouse #115-035-174) were purchased from Jackson ImmunoResearch (West Grove, PA, USA) and Trans-Blot® Turbo™ Transfer RTA kit from Bio-Rad (Hercules, CA, USA). Alexa Fluor (488 and 555)-conjugated secondary antibodies for immunofluorescence, UltraComp eBeads (#01-2222-42) for flow cytometry compensation, and mouse TNF-α ELISA kit (#88-7324-88) were purchased from Invitrogen (Carlsbad, CA, USA). The capture antibody for IFN-β ELISA (#sc-57201) was from Santa Cruz (Dallas, TX, USA), the detection antibody (#32400-1) was from PBL (Piscataway, NJ, USA), and the secondary HRP-conjugated antibody (#111-035-144) was from Jackson ImmunoResearch (West Grove, PA, USA). TMB (#ES001-500ML) was from Merck Millipore (Burlington, MA, USA). Flow cytometry anti-mouse antibodies against B220 (#103231), CD8a (#100744), CD25 (#101910), CD38 (#102707), CD44 (#103055), CD62L (#104431), CD69 (#104514), CD90.2 (#105331), FOXP3 (126404), IFN-γ (#505845), Ly-6G (#127606), MHC Class II (#107648) and NK.1.1 (#108739) were from BioLegend (San Diego, CA, USA), CD11c (#12-0114-82), CD45 (#11-0451-82), CD86 (#25-0862-80), F4/80 (#45-4801-82), KLRG1 (#11-5893-82), Ly-6C (#47-5932-82), TNF-α (#25-7321-82), IL-17A (#12-7177-81), IL-2 (#17-7021-81), and FOXP3/Transcription Factor Staining Buffer Set (#00-5523-00), and Live/Dead dye (#65-0863-14) were purchased from eBioscience (San Diego, CA, USA); CD4 (#560181), CD11b (#553312), 2.4G2 (Fc block, #553142) and Perm/Wash Buffer (#554723) were from BD Pharmingen (San Diego, CA, USA). Perm/Wash buffer (#51-2091KZ) was from BD Biosciences (Franklin Lakes, NJ, USA). Ammonium-Chloride-Potassium lysing buffer (A10492-01) was from Gibco (Waltham, MA, USA), DAPI (#D9542), PMA (#P1585), ionomycin (#I0634), and H_3_PO_4_ (#30417) were from Sigma Aldrich (St. Louis, MO, USA) and Tissue-Tek O.C.T. Compound (#4583) from Sakura (Osaka, Japan). TrueBlack (#23007) was from Biotium (San Francisco, CA, USA). Vectashield (#H-1200-10) and ImmEdge Hydrophobic Barrier PAP Pen (#VE-H-4000) were from Vector Laboratories (Burlingame, CA, USA). Pierce BCA Protein Assay Kit (#23227) was from Thermo Scientific (Waltham, MA, USA), and RayBio® C-Series Mouse Inflammation Antibody Array C1 kit (#126AAM-INF-1-8) from Raybiotech (Norcross, GA, USA). Liberase TM (#5401020001) and DNAse I (#10104159001) were from Roche (Basel, Switzerland), and Percoll from GE Healthcare (Chicago, IL, USA). Ketamine was from Richter Pharma (Wels, Austria) and xylazine was from Alfasan International (Woerden, Netherlands).

## Motor coordination and cognitive tests

### Motor coordination tests

Motor coordination was tested using a rotating rod (rotarod) and wire-hanging tests. The latency time to falling off a rotarod was measured at multiple time points during ageing from one to two years using the RotaRod apparatus (#47600, Ugo Basile S.R.L; Gemonio, Italy). The procedure had two parts: training and testing. On a training day, each mouse was given two trials at a fixed speed of 5 rpm for 4 minutes (min) and the third trial at a fixed speed of 8 rpm for 3 min. On the test day, three trials were performed at an accelerated mode from 5 rpm to 50 rpm for 300 seconds (s), and animals were allowed to rest for one hour in between each trial. For the wire-hanging test, each mouse was placed in the centre of a 3 mm thick wire and the time spent on the wire was measured (up to 300 s); an average time of three trials is shown.

### Passive avoidance test

To evaluate learning and fear-motivated memory passive avoidance test was performed. Briefly, mice were examined during a three-day trial consisting of acclimatization, training, and a testing day. On acclimatization day each mouse was given 5 min to explore both light and dark compartments, connected by an open door. On the training day, a mouse was placed in the bright compartment, and the door between the compartments opened after 30 s. Upon entering the dark compartment, a mouse received a mild foot shock of 0,3 mA. On the test day, a mouse was placed in the bright compartment, and the time needed to enter the dark compartment (latency) was measured. Mice with normal learning and memory tend to avoid entering the dark compartment during the 300 s of the test (despite their natural tendency to choose darker areas) because they remember the previous exposure to the shock.

### Novel object recognition test

The novel object recognition test was performed similarly as previously reported (32). On the first day, animals were allowed to familiarize with a plexiglass box (42 cm x 26.5 cm x 15 cm). The following day, two identical objects were placed in a box and animals were allowed to explore the area. On the testing day, one familiar object was replaced with a new one, and the time during which the animals interacted with each object was measured. Each day animals were allowed to spend 300 s in the plexiglass boxes. Preference for the novel object was calculated as discrimination index; 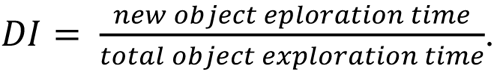

#### Tissue preparation

For immune cell characterization by flow cytometry analyses, animals were euthanized, spleens were taken out and immediately processed. For immunofluorescence and inflammation array analyses, mice were deeply anaesthetized with an intraperitoneal injection of 0.1 mg/g ketamine and 0.02 mg/g xylazine. Mice were then perfused with 60 mL of ice-cold phosphate-buffered saline (PBS). For mouse inflammation array analyses, brains, and spinal cords were removed, immediately snap-frozen in liquid nitrogen, and stored at −80°C until use. For immunofluorescence analyses, mice were perfused with PBS and fixed with 60 mL of 4% paraformaldehyde. After fixation, the brains and spinal cords were removed and postfixed overnight (O/N). The tissues were then dehydrated and kept in 30% sucrose in PBS until cryosectioning.

#### Immunofluorescence

Dehydrated tissues were embedded in O.C.T. compound on dry ice until completely frozen, and 25 μm thick coronal sections were cut on a Leica CM1850 cryostat (Leica, Wetzlar, Germany). Tissue slices were kept in a cryoprotectant solution (30% ethylene glycol, 30% glycerol, 24.4 mM phosphate buffer, pH = 7.4) until immunostaining. Tissue slices were mounted on slides, blocked, and permeabilized with 10% horse serum in 0.25% Triton X-100 in PBS and stained with primary antibodies diluted in 1% horse serum in 0.25% Triton X-100 in PBS O/N at room temperature (RT). The following day, the sections were washed with PBS and incubated with secondary antibodies (1:500) for 1 hour (h) at RT. Following washing with PBS, the sections were incubated with 500 ng/mL of DAPI for 15 min. The sections were then incubated with TrueBlack (diluted 1:20 in 70% ethanol) for 30 s to quench lipofuscin-caused autofluorescence and mounted in Vectashield. For analysis of lipofuscin, autofluorescence was measured in the green and red channels (488 and 555 nm) without prior quenching with TrueBlack. Tissue slices were imaged on Olympus IX83 fluorescent microscope (Tokyo, Japan) using 10X and 20X objectives. Z-stacking (30 stacks, 1 µm step size) was used for the visualisation of Iba1^+^ and GFAP^+^ cells. Mean fluorescence intensity (MFI), cell body area, and longest process length were analysed in ImageJ software. The brightness and contrast of representative figures were adjusted equally in Adobe Photoshop software.

#### Flow cytometry immune cell characterization

For flow cytometry analysis, cells were released from the spleen by gentle grinding. Cell suspensions were then subjected to red blood cell lysis with Ammonium-Chloride-Potassium lysing buffer for 5 min at RT. Live cell numbers were determined on Neubauer chambers upon trypan blue staining. To prevent nonspecific binding of antibodies, Fc blocking antibody (anti-CD16/CD32, clone 2.4G2) was added for 5 min prior to staining. Cells were then incubated with a mixture of primary antibodies prepared in FACS flow buffer (2% FBS in PBS) for 30 min at +4°C. After staining, cells were washed three times in FACS flow buffer and incubated with 300 nM DAPI for 5 min in the dark, washed once more, filtered through a 70 μm cell strainer, and kept on ice until analysis. Dead cells and doublets were excluded based on negativity for DAPI and forward scatter area/height gating. T and B lymphocytes were further distinguished based on positivity for T cell Thy1.2 (CD90.2) and B cell B220 (CD45R) markers. The following T cell populations were profiled: helper (CD4^+^) and cytotoxic (CD8^+^) cells, which were further subdivided into naïve populations (CD44^lo-int^CD62L^hi^), central memory (T_CM_; CD44^hi^CD62L^hi^), effector memory (T_EM_; CD44^hi^CD62L^lo^), regulatory (Treg; CD4^+^CD25^+^FOXP3^+^), and various activated subsets (CD44^+^CD38^+^, CD44^+^CD69^+^, and CD44^+^KLRG1^+^). Innate immune cells were first gated as non-T/non-B, and then further separated on neutrophils (Ly6C^+^Ly6G^+^), macrophages/monocytes (Ly6C^+^Ly6G^-^CD11b^+^F4.80^+^), conventional dendritic cells (cDc; CD11c^+^MHC-II^+^) and NK cells (NK1.1^+^). The activation status of macrophages and cDCs was monitored by MFI of CD86, CD11c, and MHC Class II. For Treg analyses, Foxp3/Transcription Factor Staining Buffer Set was used according to the manufacturer’s instructions. Briefly, cells were first stained for surface markers (CD90.2, B220, CD4, CD8, and CD25), as described above. Then the cells were subject to fixation, nuclear permeabilization, Fc blockade, and finally stained for FOXP3. For intracellular cytokine detection, cells were stimulated for 4 h with 50 ng/mL PMA and 500 ng/ml ionomycin; monensin (1:1500) was added during the whole stimulation to inhibit degranulation. The cells were first stained for surface markers (CD90.2, B220, CD4, CD8), fixed with 4% paraformaldehyde, permeabilized with Perm/Wash buffer (by manufacturer’s instructions), Fc blocked and then stained for intracellular cytokines (IL-2, IL-17, IFN-γ, and TNF-α). Flow cytometry was performed on BD FACSAria^TM^ III cytometer (BD Biosciences; Franklin Lakes, NJ, USA); data analysis was performed with FlowJo software. Absolute cell numbers of individual cell populations were calculated from the number of live cells multiplied by the percentage of indicated populations.

#### Mouse Inflammation Array

For simultaneous detection of 40 different cytokines and chemokines, brain and spinal cord lysates were analysed by a RayBio® C-Series Mouse Inflammation Antibody Array C1 kit according to the manufacturer’s instructions. Briefly, one hemisphere of the brain and spinal cord were lysed in 0.5 mL of the lysing buffer, and protein concentrations were determined using Pierce BCA Protein Assay Kit. Samples for three sets of two-year-old WT and Optn^470T^ mice were pooled together and diluted in 1X blocking buffer to obtain the final concentration of 300 μg/mL. Array membranes were blocked with 1X blocking buffer for 1 h at RT, and incubated with samples O/N at +4°C. The next day the membranes were washed three times and incubated with a cocktail of biotinylated detection antibodies O/N at +4°C. The membranes were then washed and incubated with the streptavidin-labelled antibody for 1 h at RT and developed using detection buffers on ChemiDocTM MP Imaging System (Bio-Rad; Hercules, CA, USA). Analyses were performed with ImageJ software by measuring the mean grey values of duplicates for each cytokine, normalized to the average of positive controls, and showed in the tables as means from three experimental groups ± SEM. For the graphs of selected inflammatory cytokines, data are normalized to WT.

#### Biochemical fractionation and western blot analyses

The sequential biochemical fractionation of brain tissue proteins was performed to distinguish soluble and insoluble proteins, as previously described (33). Briefly, the left hemispheres of two-year-old WT and Optn^470T^ male brains were weighed and homogenized in 5 mL/g low-salt (LS) buffer (10 mmol/L Tris, pH 7.5, 5 mmol/L ethylenediaminetetraacetic acid, 1 mmol/L dithiothreitol, 10% sucrose, and protease inhibitors) on ice. Lysates were then centrifuged at 25,000 × g for 30 min at 4°C to obtain supernatants (LS fraction), and the pellet was subjected to one more round of LS extraction and centrifugation. The remaining pellet was resuspended in 5 mL/g Triton-X (TX) buffer (LS + 1% Triton X-100 + 0.5 mol/L NaCl) and ultracentrifuged at 180,000 × g for 30 min at 4°C. Supernatants were stored (TX fraction), and the pellets were washed once more. Pellets were then homogenized in 1 mL/g sarkosyl (SARK) buffer (LS + 1% N-lauroyl-sarcosine + 0.5 mol/L NaCl) and shaken for 1 h at RT on 800 rpm, and ultracentrifuged at 180,000 × g for 30 min at RT. Supernatants were stored (SARK fraction), and the remaining pellet was resuspended in 0.5 mL/g urea buffer (7 mol/L urea, 2 mol/L thiourea, 4% 3-[(3-cholamidopro-pyl) dimethylammonio]-1-propanesulfonate (CHAPS), 30 mmol/L Tris-HCl, pH 8.5) and centrifuged at 25,000 × g for 30 min at 4°C. Supernatants were stored (urea fraction). Samples were mixed with 6X sodium dodecyl sulphate loading buffer (375 mM Tris-HCl, 12% SDS, 60% glycerol, and 0.06% bromophenol blue and 600 mM DTT), heated at 100°C for 10 min (except for the urea fraction), and separated on 12% polyacrylamide gels. Gels were transferred to nitrocellulose membranes using Trans-Blot® Turbo™ Transfer RTA kit and System on 1.3 mA for 10 min. Membranes were blocked with 3% bovine serum albumin in 0.1% Tween 20 in Tris-buffered saline and immunoblotted with the primary antibodies against TDP-43 and phospho-TDP-43 O/N 4°C. After washing, membranes were incubated with secondary antibodies and developed using a ChemiDocTM MP Imaging System (Bio-Rad; Hercules, CA, USA). Densitometric analyses were performed using ImageJ software, and data are shown as fold-change of the first (LS) WT fraction.

#### Enzyme-linked immunosorbent assay (ELISA)

TNF-α concentration in brain lysates was determined by ELISA kit from Invitrogen, following manufacturers’ instructions, while IFN-β concentration was detected as previously published (25). Briefly, 96-well microtiter plates were coated with capture antibody O/N at +4°C, washed, and subsequently blocked with blocking buffer. Brain lysates (obtained as described for cytokine array) were centrifuged at 12000 rpm for 20 min, diluted in blocking buffer in a 1:2 ratio, and incubated on plates for 2 h at RT. The plates were subsequently washed, incubated with detection antibody, washed again, and incubated with avidin-HRP conjugate (for TNF-α) or secondary HRP-conjugated antibody (for IFN-β). TMB substrate was added for 15 min, and the reaction was stopped with 8.5% H_3_PO_4_. The absorbance was read on UV/Vis spectrophotometer at 450 nm.

#### Immune cell isolation from the mouse brains

To isolate immune cells from the brain, two-month and one-year-old mice were perfused using a previously described protocol (Lee & Tansey, 2013), with minor modifications. Briefly, after perfusion with ice-cold PBS, the brains were isolated, dissected, and digested with 13 U/mL Liberase TM and 350 kU/mL DNAse I; the brain homogenates were separated using a Percoll density gradient (30%/37%/70%). Immune cells were collected from the 37%/70% Percoll interphase, left unstained or stained with 300 nM DAPI, CD45-FITC, and CD11b-APC antibodies, and analysed using BD FACSAria^TM^ III cytometer. Unstained samples were also seeded on coverslips, fixed, stained with 300 nM DAPI, and analysed on Olympus IX83 fluorescent microscope (Tokyo, Japan).

#### Statistics

Statistical analysis was performed using GraphPad Prism Software 8.0.1 (San Diego, CA, USA). For comparisons between two individual groups, Student’s t-test was used, whereas two-way ANOVA with Tukey’s *post-hoc* test was used for comparisons between multiple groups. A *p*-value of < 0.05 was considered as statistically significant.

## Results

### Optineurin insufficiency mice do not exhibit motor and cognitive deficits

To test if the optineurin insufficiency (Optn^470T^) mouse model phenocopies C-terminal optineurin truncations found in ALS patients, we assessed weight, motor functions, and cognition at several time points during a two-year period. We analysed separately males and females because of a slight male bias reported for ALS patients (1). WT males gained weight between six months and one year (13%), and then lost approximately the same amount by two years (Fig. 1A). In comparison, Optn^470T^ males weighed the same as WT mice at six months, but on average gained slightly less weight by one year (9%). For this reason, in contrast to WT mice, Optn^470T^ mice did not show significant weight loss by two years. In contrast, both WT and Optn^470T^ female mice progressively and comparably increased weight between six months and two years (Fig. 1B). To evaluate motor phenotype in WT and Optn^470T^ mice, we performed rotarod and wire-hanging tests. The average latency to fall from the rotarod in WT and Optn^470T^ males (Fig. 1C) and females (Fig. 1D) showed no significant difference at any time point. Furthermore, wire-hanging test data showed a substantial decline in motor coordination between one- and two-year-old females, but not males (Fig. 1E-F). However, there were no differences between the genotypes, corroborating the rotarod results. As ALS and FTD share common genetic backgrounds, we tested if Optn^470T^ mice show deficits in learning and fear-motivated memory. We observed no differences between WT and Optn^470T^ mice in the passive avoidance (Fig. 1G-H) and novel object recognition tests (Fig. 1I-J), demonstrating normal cognitive functions in Optn^470T^ mice. Notably, we also did not detect a different survival rate between WT and Optn^470T^ mice (data not shown). To conclude, we showed that Optn^470T^ male mice had slightly lower average weight at one year, but no overt ALS or FTD phenotype when compared to the age-matched WT mice.

**Figure 1.**
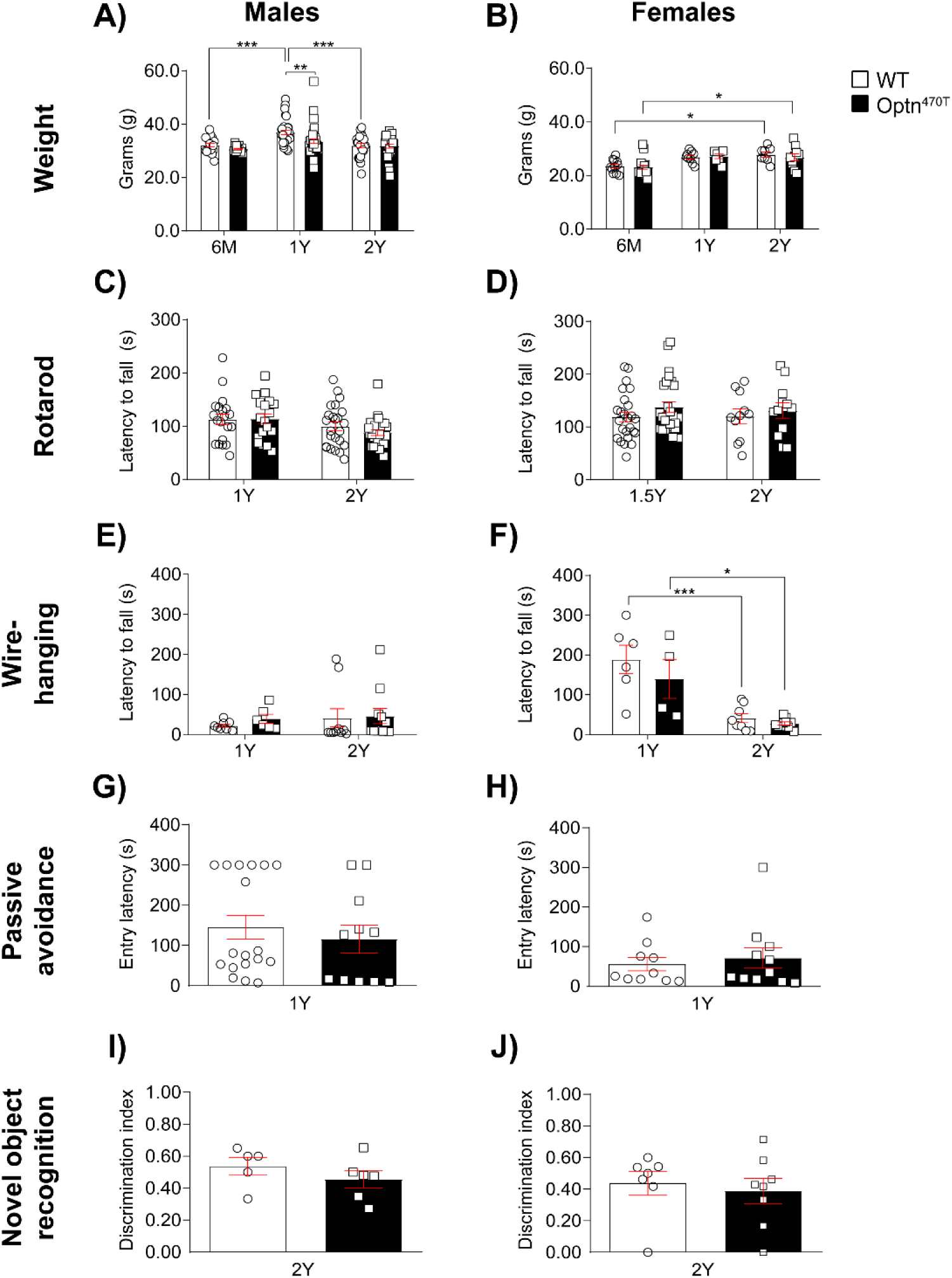
Optineurin insufficiency did not affect motor coordination and cognition. Bar diagrams show body weight (in grams) for individual males at the age of six months (20 WT, 17 Optn^470T^), one year (35 WT, 39 Optn^470T^), and two years (26 WT, 29 Optn^470T^) **(A)**, and females at the age of six months (11 WT, 16 Optn^470T^), one year (9 WT, 7 Optn^470T^) and two years (8 WT, 10 Optn^470T^) **(B)**. Motor coordination on the rotarod measured as the average latency to fall (in seconds) is shown for males **(C)** at the age of one year (20 WT, 18 Optn^470T^) and two years (24 WT, 28 Optn^470T^), and females **(D)** at the age of one and a half years (25 WT, 26 Optn^470T^), and two years (11 WT, 12 Optn^470T^); in the wire-hanging test for males **(E)** at the age of one year (8 WT, 6 Optn^470T^), and two years (10 WT, 11 Optn^470T^), and females **(F)** at the age of one year (6 WT, 4 Optn^470T^), and two years (8 WT, 10 Optn^470T^). Evaluation of fear-motivated memory is shown as entry latency (in seconds) in the passive avoidance test for males (18 WT, 11 Optn^470T^) **(G)** and females (10 WT, 11 Optn^470T^) **(H)**. Discrimination index in novel object recognition test is shown for males (5 WT, 6 Optn^470T^) **(I)**, and females (7 WT, 8 Optn^470T^) **(J)** at two years. Data are presented as means ± SEM from the indicated number of mice and analysed by two-way ANOVA (A-F) and Student’s t-test (G-J): * p < 0.05, ** p < 0.01, *** p < 0.001.

### Ageing did not precipitate neuropathology in mice with optineurin insufficiency

To test if ageing precipitates the development of ALS-like neuropathology in mice carrying the optineurin truncation, we analysed the lumbar spinal cords and motor cortex of one- and two-year-old mice. We observed a trend towards increased astrocyte activation between one- and two-year-old WT male mice in the lumbar spinal cords (Fig. 2A-B). Optn^470T^ mice exhibited a similar level of astrocyte activation as WT mice. The glial fibrillary acidic protein (GFAP) signal was not quantified in the motor cortex since we detected very few astrocytes per field (data not shown). During ageing, we observed increased microglial activation in the lumbar spinal cords of WT animals, visible as larger cell body area and shortening of processes (Fig. 2C-E) without substantial difference in Iba1 positivity (Fig. 2F). Increased activation was absent from the motor cortex between one and two years of age (Suppl. Fig. 1A-D). There was no difference in microglial activation between the genotypes (Fig. 2D-F). Furthermore, no difference between the genotypes or between ages was found in motor neuron numbers in the lumbar spinal cords (Fig. 2G-H) or motor cortex (Suppl. Fig. 1E-F). Next, we analysed the MFI of lipofuscin (measured as autofluorescence), which was shown to accumulate during ageing (35). We observed a significant increase in the lipofuscin MFI signal (∼60%) in the lumbar spinal cords and the motor cortex between one- and two-year-old mice but without any difference between the genotypes (Fig. 2I-J, Suppl. Fig. 1G-H). Lastly, we checked for the presence of TDP-43 protein aggregation. Because of the reportedly different preferential staining of cytoplasmic and nuclear TDP-43 by antibodies targeting the C- and N-terminus of TDP-43, respectively (26), we used both antibodies. None of the anti-TDP-43 antibodies detected cytoplasmic aggregates (Suppl. Fig. 1J-L), whereas the N-terminal antibody clearly showed predominant nuclear localisation of TDP-43 (Suppl. Fig. 1J-K). To further strengthen this result we performed staining for phospho-TDP-43, because pathological TDP-43 aggregates are phosphorylated (36). We found no phosphorylated TDP-43 aggregates in the lumbar spinal cords or motor cortex of Optn^470T^ mice at any age tested (Fig. 2K and Suppl. Fig. 1I). Hyperphosphorylated TDP-43 aggregates are detergent-insoluble, so we tested TDP-43 solubility in brain homogenates of two-year-old mice by biochemical fractionation. TDP-43 was found only in detergent-soluble LS and TX fractions (Fig. 2L-M), whereas phospho-TDP-43 was present in insoluble SARK and urea fractions (Fig. 2L and N). Nevertheless, no signs of differential TDP-43 insolubility or hyperphosphorylation were found between the genotypes. In conclusion, we observed marked signs of ageing in both WT and Optn^470T^ mice, but without ALS-like neuropathology in the latter.

**Figure 2.**
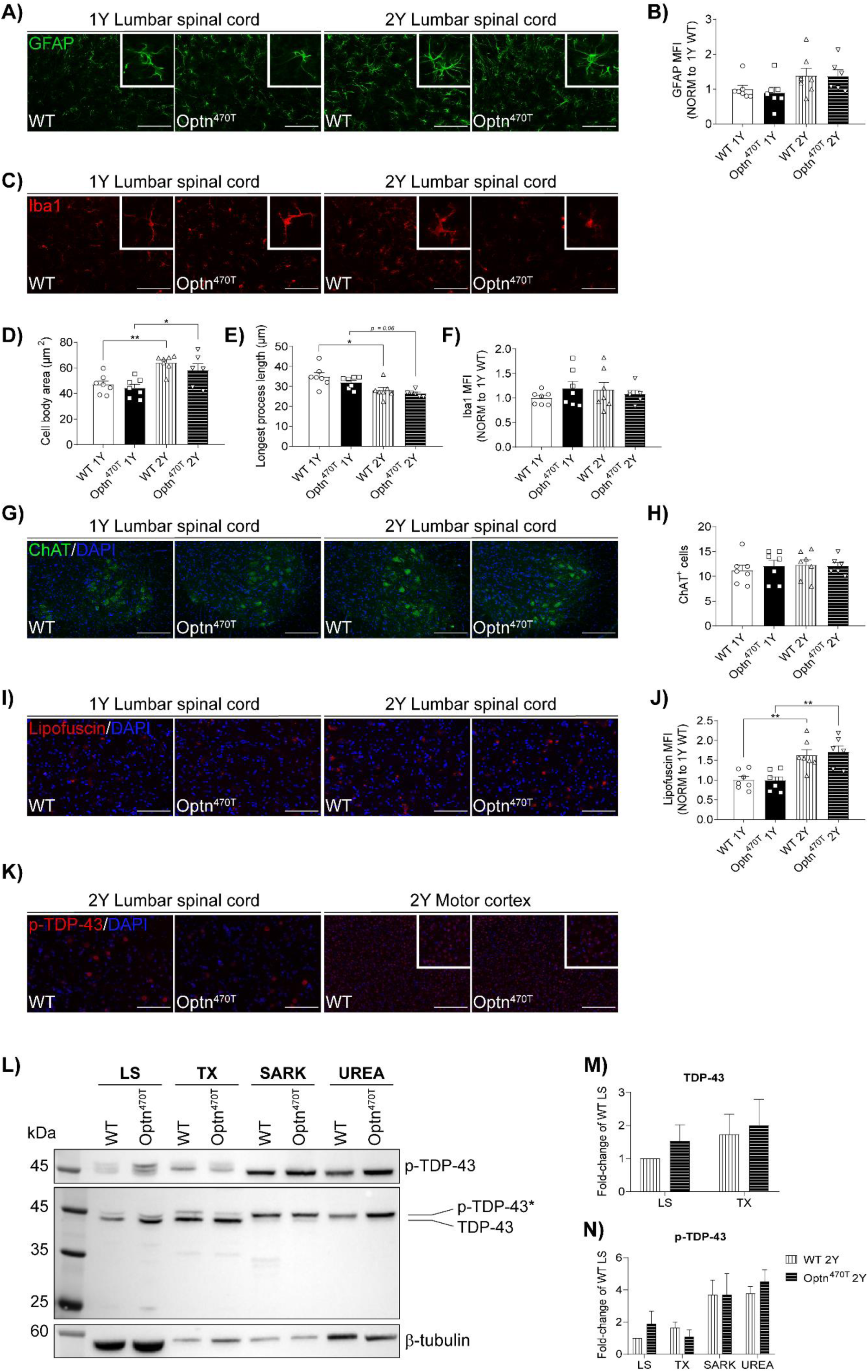
Ageing did not induce ALS-like neuropathology in Optn^470T^ mice. Lumbar spinal cord sections from one- and two-year-old male mice were stained for GFAP **(A)**, Iba1 **(C)**, ChAT **(G)**, and phospho-TDP-43 **(K)** or were left unstained for lipofuscin visualization **(I)**; nuclei were identified by DAPI staining. MFI of GFAP **(B)** and Iba1 **(F)**, microglial cell body area **(D),** longest process length **(E)**, ChAT^+^ cell number **(H)**, and lipofuscin **(J)** are shown. Data are presented as means ± SEM from 6-7 one-year- and two-year-old WT and Optn^470T^ mice analysed by two-way ANOVA: * p < 0.05, ** p < 0.01. The scale bar is 100 μm (A, C, I, and K) and 200 μm (G and K). The brains of individual two-year-old WT and Optn^470T^ mice were homogenized, biochemically fractionated, and immunoblotted for TDP-43, p-TDP-43, and β-tubulin **(L);** remark: β-tubulin is relevant as a loading control only for low-salt (LS) fraction. Densitometric analysis of TDP-43 **(M)** and phospho-TDP-43 **(N)** is shown as a fold-change of WT in the LS fraction; TX = Triton X fraction; SARK = sarkosyl fraction; data represent means ± SEM from three mice per genotype.

### Cytokine and chemokine expression is comparable in the aged brains and lumbar spinal cords of Optn^470T^ and WT mice

Upon finding no neuropathological differences in the CNS between Optn^470T^ and WT mice during ageing, we assessed potential functional outcomes in the inflammatory status by analysing the levels of 40 different pro- and anti-inflammatory cytokines, chemokines, and growth factors in the brain and spinal cord lysates in two-year-old male mice by a protein array. Cytokine expression in the brain and spinal cord lysates of WT and Optn^470T^ mice was similar (Suppl. Fig. 2A). In the brains the highest expression was found for fractalkine, a chemokine by which neurons suppress microglial activation, chemoattractants and/or growth factors LIX, CCL2, and M-CSF, and IL-1α and IL-4, a pro- and anti-inflammatory cytokine, respectively (Fig. 3A, Suppl. Fig. 2B), but there were no differences between Optn^470T^ and WT mice. The only difference between the genotypes was a higher level of I-TAC (CXCL11; a chemoattractant for activated T cells) in Optn^470T^ compared to WT brains (Fig. 3A). Since we detected a low expression level of a major proinflammatory cytokine tumour necrosis factor (TNF)-α in the protein array, we also performed an ELISA. We obtained a similar result – a low level of TNF-α, indistinguishable between the genotypes (Fig. 3B). IFN-β was either expressed at low level or was below the level of detection by ELISA, but there was also no difference between WT and Optn^470T^ mice (Fig. 3C). Compared to brains, spinal cords had a higher level of XCL1 in Optn^470T^ mice, with no other significant differences (Fig. 3D and Suppl. Fig. 2C). Overall, we observed no changes in major pro- and anti-inflammatory factors in Optn^470T^ and WT brains and spinal cords, with minor differences detected in cytokines I-TAC and XCL-1.

**Figure 3.**
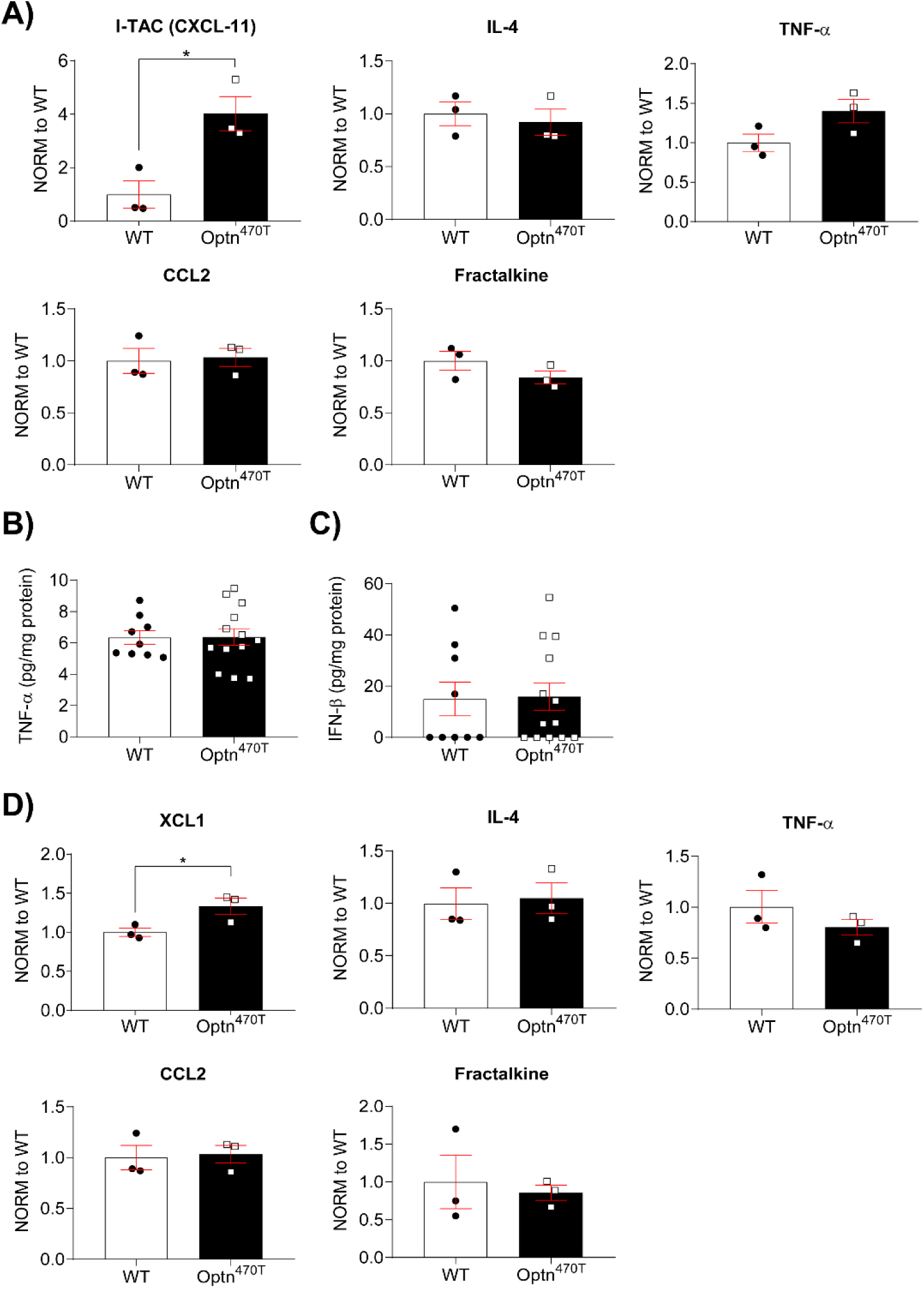
Similar inflammatory profiles in the brains and spinal cords of aged Optn^470T^ and WT mice. RayBio® C-Series Mouse Inflammation Antibody Array C1 was performed on brain and spinal cord lysates from two-year-old male WT and Optn^470T^ mice. Selected pro- and anti-inflammatory cytokines in the brain **(A)** and spinal cord **(D)** from WT and Optn^470T^ mice are shown normalized to WT. Three independent experiments were performed from the following groups of mice: 1) 4 WT and 5 Optn^470T^; 2)1 WT, 5 Optn^470T^; 3) 4 WT, 3 Optn^470T^. Brain lysates from 9 two-year-old male WT and 13 Optn^470T^ mice were assayed by ELISA for TNF-α **(B)** and IFN-β **(C)**. The data were analysed by Student’s t-test: * p<0.05.

### Optineurin insufficiency did not affect ageing-induced T cell phenotype

To determine if optineurin insufficiency affects T cell subsets and their activation during ageing, we profiled splenocytes of one- and two-year-old WT and Optn^470T^ mice by flow cytometry (gated as shown in Fig. 4A). We separately analysed males (Fig. 4 and Suppl. Fig. 3) and females (Suppl. Fig. 4). Of note, splenocyte numbers were comparable between the genotypes at both one and two years, with higher mouse-to-mouse variability seen in older mice (Suppl. Fig. 3A). B cell frequencies were similar in WT and Optn^470T^ mice during ageing (Suppl. Fig 3B), but absolute B cell numbers slightly decreased at two years in WT mice (Suppl. Fig 3C). T cell frequencies remained unchanged over time and between genotypes (Fig. 4B), with a trend of decreased absolute T cell numbers in WT at two years (Fig. 4F). The percentages of CD4^+^ and CD8^+^ T cells were comparable in one- and two-year-old WT and Optn^470T^ mice (Fig. 4C). A slight increase in CD8^+^ T cells frequency was observed from one to two years in Optn^470T^ mice, but this did not result in perturbation of CD4^+^/CD8^+^ ratio between the genotypes (Suppl. Fig 3D). The absolute numbers of both CD8^+^ and CD4^+^ T cells significantly declined in two-compared to one-year-old WT males, while the numbers in Optn^470T^ mice remained the same (Fig. 4G).

**Figure 4.**
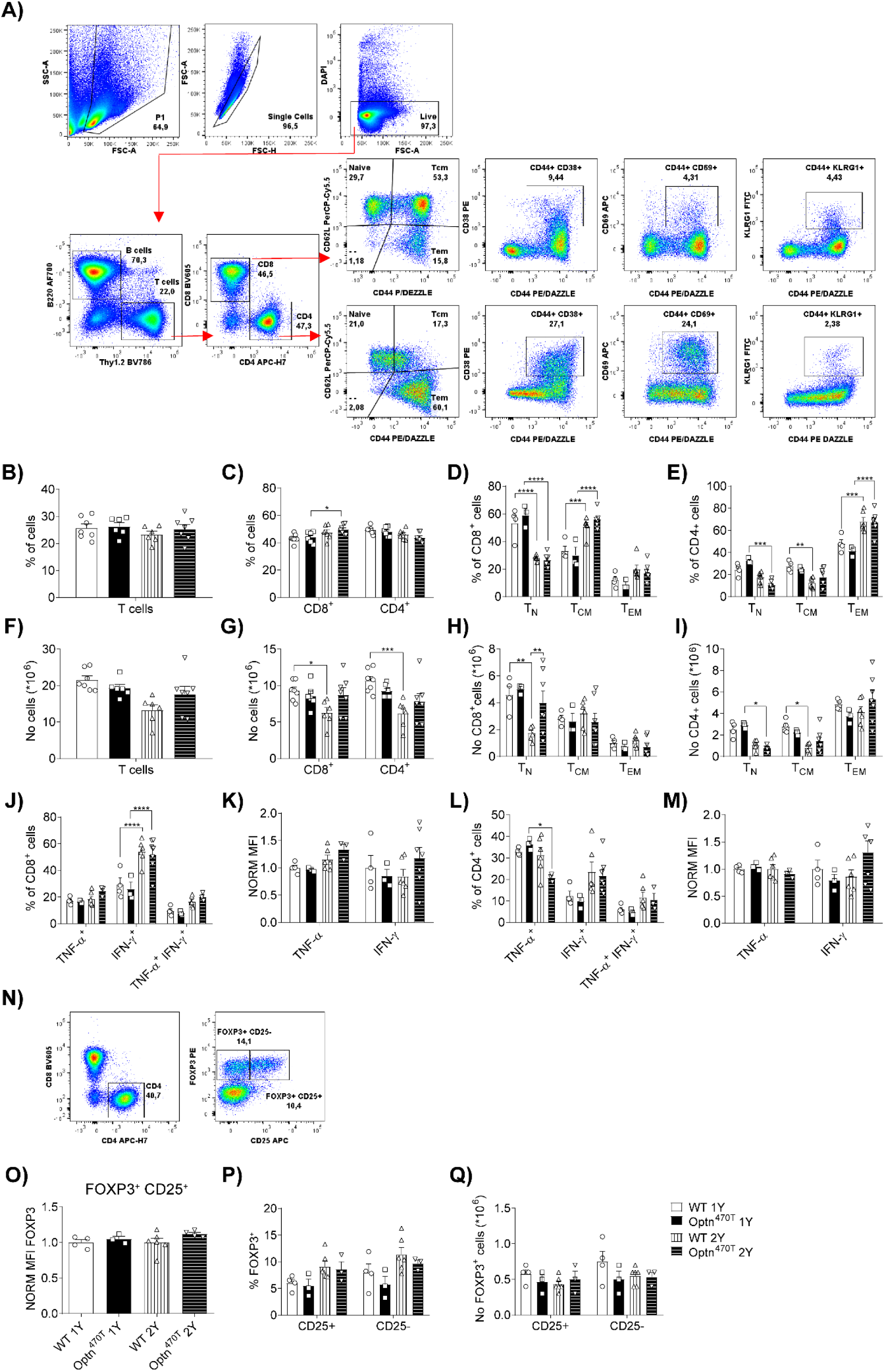
T cell subsets in aged Optn^470T^ were comparable to WT male mice. The gating strategy in mouse spleens is shown for a representative two-year-old mouse **(A)**. Population frequency (%) and absolute cell numbers (No) are shown for one- and two-year-old WT and Optn^470T^ as follows: frequency of T cells **(B)**, CD8^+^ and CD4^+^ T cells **(C),** CD8^+^ naïve (T_N_), central memory (T_CM_) and effector memory (T_EM_) **(D)**, and CD4^+^ T_N_, T_CM_ and T_EM_ **(E)**; numbers of T cells **(F)**, CD8^+^ and CD4^+^ T cells **(G)**, CD8^+^ T_N_, T_CM_ and T_EM_ **(H)**, and CD4^+^ T_N_, T_CM_ and T_EM_ **(I)**. Staining for the indicated cytokines upon PMA/ionomycin stimulation is shown as frequencies and MFI in CD8^+^ (**J-K)** and CD4^+^ **(L-M)** T cells. The gating strategy for Treg cells in mouse spleens for representative two-year-old mice is shown. Functional Tregs were gated as CD4^+^FOXP3^+^CD25^+^, and non-functional as CD4^+^FOXP3^+^ CD25^-^ cells **(N)**. The graph shows the MFI of FOXP3 in Optn^470T^ normalized to one- and two-year-old WT mice in functional Tregs **(O)**. Treg subsets are shown as frequency **(P)** and numbers **(Q)**. The data from 6-7 one-year- and two-year-old WT and Optn^470T^ mice are shown as means ± SEM and analysed by two-way ANOVA: * p<0.05, ** p<0.01, ***p<0.001, ****p<0.0001.

Several ageing-related phenotypes were observed between one- and two-year-old WT mice including a decreased frequency and/or numbers of naïve and increased percentage of memory CD4^+^ and CD8^+^ T cells (Fig. 4D-E and H-I). Of note, CD8^+^ T cells had a greater increase in the percentage of T_CM_ and CD4^+^ T cells in T_EM_ (Fig. 4D-E). Notably, though, Optn^470T^ T cells exhibited the same ageing phenotype as WT mice. We also analysed various activated T cell subsets and demonstrated that Optn^470T^ mice had a slightly higher number of CD8^+^CD44^+^CD38^+^ T cells (Suppl. Fig. 3F) and a higher percentage and number of CD4^+^CD44^+^CD38^+^ T cells (Suppl. Fig. 3G-H). The percentage of CD4^+^CD69^+^CD44^+^ T cells was significantly increased in both WT and Optn^470T^ mice at two years compared to one year, but a significant increase in absolute CD4^+^CD69^+^CD44^+^ T cell numbers was observed only in Optn^470T^ mice (Suppl. Fig. 3G-H). These slight differences in activated CD4^+^ or CD8^+^ T cell subsets in Optn^470T^ mice did not translate into a higher frequency of TNF-α-, IFN-γ- (Fig. 4J and L), IL-2- or IL-17-secreting cells (Suppl. Fig. 3I and K) upon PMA/ionomycin stimulation. Nevertheless, we expectedly observed substantially increased levels of IFN-γ^+^ CD8^+^ T cells between two- and one-year-old mice in both genotypes (Fig. 4J). A decreased percentage of TNF-α-secreting CD4^+^ T cells was observed in two-year-old Optn^470T^ mice (Fig. 4L). Notably, ageing did not increase cytokine secretion per cell as there was no change in MFI for any of the cytokines tested (Fig. 4K and M, and Suppl. Fig. 3J and L). We also checked regulatory CD4^+^ T cells (Treg; gated in Fig. 4N) because of the reported decrease of Tregs in ALS models and patients’ blood (6). Tregs were subdivided into functional (CD4^+^FOXP3^+^CD25^+^) and non-functional (CD4^+^FOXP3^+^CD25^-^), as previously reported in aged mice (37), The percentages, numbers, and FOXP3 MFI of Tregs were comparable between WT and Optn^470T^ mice and did not substantially change between one and two years (Fig. 4O-Q).

The same analyses performed in female mice demonstrated overall similar findings as male mice (Suppl. Fig. 4). The differences included a slightly more prominent decrease in B cell numbers at two years compared to one year (Suppl. Fig. 4B-C) and an increase in the percentage and/or numbers of CD8^+^ and CD4^+^ T_EM_ in both WT and Optn^470T^ females at two years (Suppl. Fig. 4G and K and 4H and L). Furthermore, while Treg absolute numbers were the same as in males, both WT and Optn^470T^ two-year-old females showed an increase in the percentage of non-functional (CD25^-^) Tregs compared to one-year-old females (Suppl. Fig. 4V-W). Notably, Tregs in two-year-old Optn^470T^ females exhibited slightly lower FOXP3 MFI (Suppl. Fig. 4U). In summary, we showed that both male and female two-year-old WT and Optn^470T^ mice showed T cell differences typical for ageing (decrease of naïve and increase of memory cells; increased IFN-γ secretion), but otherwise we detected negligible differences between the genotypes.

### Optineurin insufficiency caused increased activation of cDC in two-year-old male mice without affecting the numbers and percentages of innate immune cell subsets

To further characterize inflammageing in Optn^470T^ mice we analysed innate immune cells and their activation status in the spleens of old mice (gating is shown in Fig. 5A). No differences were found in the numbers or frequencies of non-T/non-B cells (Fig. 5B and E), cDC, macrophages (Fig. 5C and F) and, neutrophils and NK cells (Fig. 5D and G) between WT and Optn^470T^ one- and two-year-old male mice. As previously described in ageing process, an increase in the percentage of neutrophils was observed between one and two years, which was significant only in WT mice due to higher Optn^470T^ mouse-to-mouse variability (Fig. 5D). However, the total numbers of neutrophils did not increase over time in any of the genotypes (Fig. 5G). Notably, we observed increased activation of cDC and macrophages from the Optn^470T^ mice at two years, which was visible by higher expression of CD86 and/or MHC-II activation markers (Fig. 5H-I). In comparison to males, two-year-old females had higher percentages of total non-T/non-B cells (Suppl. Fig. 5A), which was mostly caused by an increase in macrophages (Suppl. Fig. 5E), without significant changes in other populations (Suppl. Fig. 5B-F). However, in contrast to males, Optn^470T^ females did not exhibit higher cDC and macrophage activation at two years (Suppl. Fig. 5G-H). In conclusion, ageing led to an increased frequency of neutrophils at two years in male but not female mice. Furthermore, optineurin insufficiency caused increased activation of cDC and macrophages in males but not in females.

**Figure 5.**
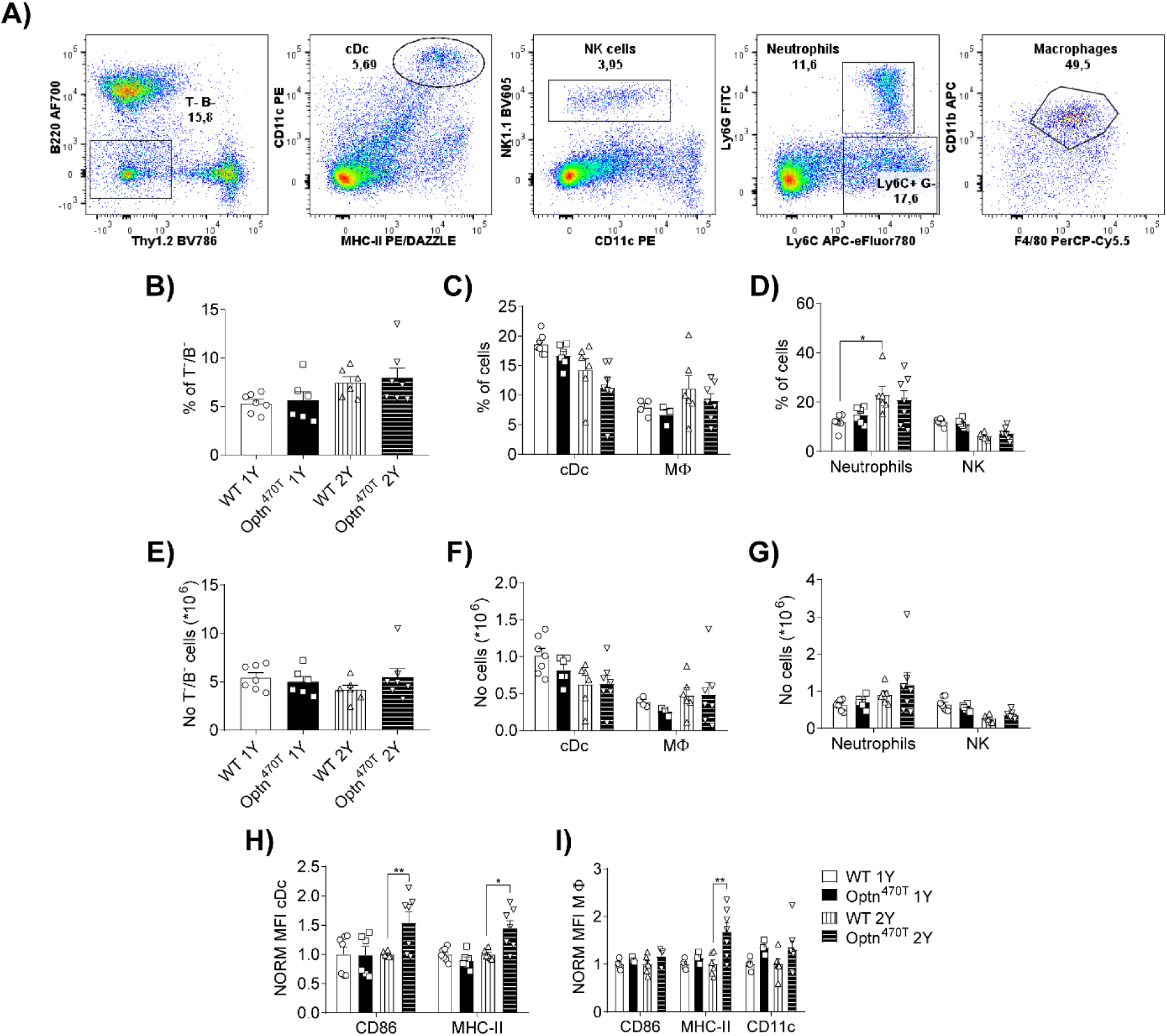
Optineurin insufficiency showed increased activation of cDC in two-year-old male mice. The gating strategy for innate immune cells in mouse spleens is shown for a representative two-year-old mouse **(A)**. Population frequency (%) and absolute cell numbers (No) are shown as following: frequency of non-T/non-B cells **(B)**, conventional dendritic cells (cDc) and macrophages (MΦ) **(C)**, and neutrophils and NK cells **(D)**; numbers for non-T/non- B **(E),** cDc and MΦ **(F)**, and neutrophils and NK cells **(G)**. The MFI for indicated activation markers normalized to one- and two-year-old WT and Optn^470T^ mice are shown for cDc **(H)**, and MΦ **(I)**. The data from 6-7 one-year- and two-year-old WT and Optn^470T^ mice are shown as means ± SEM and analysed by two-way ANOVA: * p<0.05, ** p<0.01.

## Discussion

The purpose of the study was to test if ageing would provide a sufficient second hit to elicit neuropathology in Optn^470T^ mice. Our previous studies reported no overt phenotype in unmanipulated young adult Optn^470T^ mice (31,24). Lack of phenotype was also reported for Optn^D477N^ mice, designed to mimic ALS/FTD-linked pathogenic point mutation in the ubiquitin-binding domain (17). In contrast, a disbalanced inflammatory signalling in various myeloid cells derived from these models was reported in all of these studies (17,31,24). Thus, we hypothesized that ageing, as a major risk factor for neurodegenerative diseases, would aggravate the immune system senescence and/or inflammageing in Optn^470T^ mice and trigger neurodegeneration. We observed that aged Optn^470T^ male, but not female mice showed slightly lower average weight compared to WT mice at one year, although this difference was lost at a two-year time-point. Notably, in both males and females, we failed to find differences in motor (rotarod and wire-hanging) and cognitive (novel object recognition and passive avoidance) tests between WT and Optn^470T^ mice, suggesting that optineurin insufficiency was unable to induce an ALS/FTD-like phenotype. Therefore, these results showed that Optn^470T^ mice were similar to optineurin-deficient Optn^-/-^ mice made by several individual groups, which also showed no obvious neurological phenotype during ageing (23,38,21,22), except for a mildly decreased frequency of hind leg usage reported by Ito *et al.* (15). The latter report also showed spinal cord dysmyelination, which was contested in a follow-up study (38). Of note, a decrease in motor functions was found in 10-week-old mice that received a stereotactic injection of lentiviral vector encoding for the ALS/FTD patient OPTN^E478G^ mutation directly into the motor cortex (18). However, considering the comparable findings in Optn^-/-^ and Optn^470T^ mice, the (patho)physiologic relevance of such a model is unclear because the observed phenotype could be an artefact of OPTN^E478G^ protein overexpression or triggered by some unknown dominant-negative effect of OPTN^E478G^. It is of note though that the above-mentioned Optn^D477N^ mouse model, which, unlike the lentiviral OPTN^E478G^ model, expresses physiological levels of ubiquitin-binding optineurin mutation, did not exhibit overt ALS symptoms up to one year of age, although no detailed neurological or neuropathological analysis was reported (17). Overall, we can conclude that similar to previously published Optn^-/-^ models (23,38,21,22), ageing up to two years was insufficient to elicit ALS/FTD phenotype in a ubiquitin-binding-deficient Optn^470T^ mouse model.

Since motor neuron loss and glial activation are reported to precede the symptom onset (7), we performed extensive neuropathological analyses of the lumbar spinal cords and motor cortex of ageing WT and Optn^470T^ mice. Given that men are slightly more affected by ALS than women (1), for these analyses we focused on male mice. Analysis of the one- and two-year-old spinal cords revealed a significant increase in several hallmarks of ageing, including higher microglial activation and lipofuscin accumulation. We also observed a trend toward increased astrocyte activation. Lipofuscin was also significantly increased in two-compared to the one-year-old motor cortex, whereas other ageing markers were not increased or showed only a trend toward an increase. This was likely because we compared two- to one-year-old mice, rather than to younger adults. Of note, flow-cytometry analysis of lipofuscin already showed a several-fold increase between two-month- and one-year-old mice, corroborating this finding by showing that ageing phenotype is already evident at one year of age (Suppl. Fig. 6A-B). Nevertheless, we found no differences between the genotypes. Moreover, neuronal numbers were preserved between one and two years of age in both the lumbar spinal cords and the motor cortex of WT and Optn^470T^ mice. The preserved motor neuron numbers in Optn^470T^ mice compared to WT aged mice thus corroborated the previous findings of no neuronal loss in Optn^-/-^ mice (15,38). In contrast to our study and two studies in Optn^-/-^ mice, Kurashige *et al.* reported a significantly decreased lumbar spinal but not cortical motor neuron numbers from six months to two years of age in Optn^-/-^ compared to WT mice (21). It is notable though that this study reported considerable mouse-to-mouse variability and preserved motor function, with the latter being similar to our study. Another discrepancy that we found with Kurashige *et al.* was in TDP-43. The cytoplasmic phosphorylated TDP-43 aggregates and the depletion of nuclear TDP-43 are hallmarks of ALS and FTD (4,5). Studies in patients with a heterozygous E478G and homozygous Q398X optineurin mutation have also shown TDP-43-positive neuronal cytoplasmic inclusions in spinal and bulbar motor neurons (11,39). However, in the aged Optn^470T^ mice we found no TDP-43 aggregates in neither lumbar spinal cords nor the motor cortex, whereas Kurashige *et al.* reported TDP-43 mislocalization and aggregation in the lumbar spinal cords. Therefore, to directly compare our results with that study, we performed staining with the same antibody against TDP-43 (Proteintech, #10782-2-AP; Suppl. Fig. 1L) and noticed an unusual diffuse cytoplasmic staining in both WT and Optn^470T^ mice, which is atypical for mostly nuclear TDP-43 in WT mice. To avoid ambiguity, we also used an antibody against phosphorylated TDP-43 that is more specific to aggregated TDP-43, and showed the absence of phosphorylated TDP-43 aggregates, thus making a strong case against the presence of TDP-43 aggregates in Optn^470T^ mice. The latter was also corroborated by biochemical fractionation of the whole brain. In conclusion, the findings in Optn^470T^ mice go in line with previously published results in which optineurin deficiency was insufficient to induce ALS/FTD-like neuropathology in aged mice.

In addition to analysing morphological signs of microglial and astrocyte activation, we also analysed their function by measuring cytokine, chemokine, and growth factor expression in the brain and spinal cord lysates of aged Optn^470T^ mice. This was of particular interest because optineurin has been reported to affect inflammatory signalling in various primary myeloid cells including microglia (17,31,40,23,24). Moreover, mildly increased levels (5-15%) of several proinflammatory cytokines (TNF-α, IL-1α, IL-1β, IL-2, IL-12, and IFN-γ) were detected in the lumbar spinal cords but not in the brains of Optn^-/-^ mice (15), whereas a significantly higher level of IL-1β was present in the brains with lentivirus-mediated overexpression of OPTN^E478G^ applied to the motor cortex (18). In contrast to these reports, in a protein array, we found no differences in major pro- and anti-inflammatory factors in the brains and the lumbar spinal cords in two-year-old mice. The most highly expressed factors were fractalkine, LIX, CCL2, M-CSF, IL-4, and IL-1α, which showed no differences between Optn^470T^ and WT mice. A major proinflammatory cytokine TNF-α was expressed at a low level, so we tested it further by an ELISA, where we confirmed a low level of expression and no difference between the genotypes. Small differences in Optn^470T^ male mice were found for I-TAC in the brains and XCL1 in the spinal cords, which were expressed at low levels, but could not link them to any phenotype. We also separately tested IFN-β since we previously found differences in its production by lipopolysaccharide-stimulated neonatal microglia (24). However, the levels of IFN-β were indistinguishable between the unmanipulated two-year-old brains of WT and Optn^470T^ mice. Our results showed that ageing is by itself insufficient to uncover the difference in inflammatory factor expression in the CNS between these genotypes. This was different to a slight proinflammatory bias in Optn^-/-^ mice reported by Ito *et al.* (15), but the reason behind this is unclear. It is possible that they analysed a different time-point or sex, but we cannot comment on that since the details were not reported. In our study, we analysed the lysates of 9-13 pooled male mice from three individual experiments and confirmed the TNF-α result by an ELISA, in which we analysed individual samples.

In our attempt to analyse the Percoll-enriched immune cells in the CNS by flow cytometry, we encountered a technical problem because of a large accumulation of lipofuscin, which led to a strong autofluorescence in several channels already at one year of age (Suppl. Fig. 6A-B), precluding further immunophenotyping of aged brains and the spinal cords. The autofluorescence was also confirmed by fluorescence microscopy, which showed lipofuscin accumulation in one-year-but not two-month-old mice (Suppl. Fig. 6C). For this reason, we immunophenotyped only the peripheral immune system. This was of considerable importance as well because various perturbations in the peripheral innate and adaptive immune cells were reported both in ALS and FTD (10,6,7). Importantly, despite the reported roles of optineurin in immune signalling, detailed immunophenotyping has not yet been performed in any of the optineurin mouse models. Moharir *et al.* just analysed the ratio of CD4^+^ and CD8^+^ T cells in the blood of one-year-old mice, reporting a slight reduction (22). Here, we have found age-related changes such as an increased percentage of memory and decreased percentage of naïve T cells, with an increased percentage of IFN-γ-secreting CD8^+^ T cells upon restimulation, but without any substantial differences between the WT and Optn^470T^ mice. A difference was found in activation markers of cDC and macrophages, which were increased in two-year-old Optn^470T^ male but not female mice. A somewhat opposite finding was reported for the conditional CD11c-specific optineurin knock-out model (41). Young adult CD11c-specific optineurin knock-out mice had lower expression of activation markers in cDCs, a higher percentage of Tregs, and a lower percentage of IFN-γ^+^ and IL-17^+^ CD4^+^ T cells upon stimulation. Because of this discrepancy, which could also be age-related, further investigations are necessary to understand the importance of optineurin in dendritic and T cell activation. It is of note though that detailed immune cell characterization performed in C9ORF72^-/-^ mouse model showed a higher CD86 expression in splenic cDCs in 5-month-old mice (42), which is in line with results in two-year-old Optn^470T^ mice. In conclusion, we found no evidence of advanced immunosenescence and/or inflammageing in the Optn^470T^ model and unlike the previous report in blood (22), we found an expected CD4^+^/CD8^+^ T cell ratio in spleens at both one and two years of age.

Taken together, a double hit of optineurin insufficiency and ageing was not sufficient for the development of ALS and/or FTD pathogenesis. This is reminiscent of recently analysed TBK1 haploinsufficiency (*Tbk1^+/-^*) and TBK1 ALS-patient missense mutation (G217R and R228H)-carrying mice (43,44). These models are of relevance to our research since TBK1 and optineurin interact in both inflammatory signalling pathways and autophagy (19,45). None of these models developed ALS-like neuropathology upon ageing (43,44), consistent with our results. However, when crossed with SOD1^G93A^ transgenic mice, TBK1 insufficiency was shown to be detrimental in the early and beneficial in the late stage of the disease (46). Therefore, two ALS-linked genetic mutations and ageing were necessary to elicit neuropathology in TBK1 models. This demonstrates the challenges of developing relevant ALS/FTD mouse models. The gold standard model in ALS research is a transgenic mouse expressing multiple copies of a patient SOD1^G93A^ mutation, which develops ALS-like pathology, neuroinflammation, and precociously dies at 5-6 months of age (47,48). However, this model has manyfold overexpressed SOD1 and is not representative of > 95% ALS cases, which are not SOD1^G93A^ carriers and exhibit TDP-43 rather than SOD1 aggregation. In contrast to that model, various new models have been generated, but most of them fail to phenocopy key aspects of human disease. This is likely because ALS is a multistep process with a complex genetic and environmental interplay (49). Notably, the number of steps required for disease onset is reduced in patients carrying genetic mutations (49). This is similar to the findings in TBK1 mouse models, which showed an acceleration of disease onset in SOD1 mice by TBK1 insufficiency (Brenner et al., 2019). Therefore, we would argue that optineurin models that closely mimic individual ALS and FTD patient mutations are still of major relevance, but that additional environmental and genetic challenges should be carefully selected.

## Abbreviations

ALS: amyotrophic lateral sclerosis
cDC: conventional dendritic cells
FTD: frontotemporal dementia
GFAP: glial fibrillary acidic protein
IFN: interferon
MFI: mean fluorescence intensity
NK cells: natural killer cells
O/N: overnight OPTN-optineurin
PBS: phosphate-buffered saline
RT: room temperature
SOD1: superoxide dismutase 1
TBK1: TANK-binding kinase 1
TDP-43: TAR DNA-binding protein 43
TNF: tumour necrosis factor
WT: wild-type

## Acknowledgments

We kindly thank Professor Tanja Celic at the Medical Faculty of Rijeka for providing access to their cryostat (Leica CM1850) and Marija Mrsic and Snjezana Simac Vorkapic for expert technical assistance. We also thank our lab member Marta Kolaric for her helpful discussions.

## Authors Contributions

NM and JP contributed equally to this work. Study design: IM, NM, and JP; experimental design, performance, and analyses: NM, JP, IM, AM and RC; funding acquisition: IM and BR; methodology: NM, JP, IM, AM, RC, HJ, ZM, BR, JN and MD; manuscript writing: IM, NM, and JP; manuscript editing: IM, NM, JP, RC, ZM, AM, BR, JN, MD and HJ. All authors have read and agreed to the published version of the manuscript.

## Funding

This work was funded by the Croatian Science Foundation IP-2018-01-8563 and the University of Rijeka grant 18-211-1369 to IM. The equipment was purchased from the European Regional Development Fund (ERDF) grant for the project “Research Infrastructure for Cam-pus-based Laboratories at the University of Rijeka”. NM is supported by DOK-2018-09-7739 and JP by DOK-2020-01-8703 grants to IM. BR and JN were supported by the Slovenian Research Agency (grant numbers N3-0141, J3-9263, J3-8201, J3-3065, and P4-0127).

## Ethical Approval

This study has been approved by the Ethics Committee at the Medical Faculty (University of Rijeka; Approval Code: 2170-24-18-3) and the Ministry of Agriculture of the Republic of Croatia (Approval Code: 525-10/0255-19-4).

## Conflicts of Interest

The authors declare no conflict of interest.

## Figure legends

**Suppl. Figure 1. Ageing did not induce ALS-like neuropathology in Optn^470T^ mice, continued.** Motor cortex or lumbar spinal cord sections of one- and/or two-year-old mice were stained for Iba1 **(A)** and NeuN **(E),** phospho-TDP-43 **(I)**, N-terminal TDP-43 **(J-K)**, C-terminal TDP-43 **(L)** or were left unstained for lipofuscin visualization **(G)**; nuclei were identified by DAPI staining. MFI for Iba1 **(B)**, microglial cell body area **(C)**, longest process length **(D)**, NeuN^+^ cell number **(F)**, and lipofuscin MFI **(H)** are shown. Data are presented as means ± SEM from 3-7 one-year- and two-year-old WT and Optn^470T^ mice and analysed by two-way ANOVA: **p<0.01, ***p<0.001. The scale bar is 100 μm (I, J, and L) and 200 μm (A, E, G, I, and K).

**Suppl. Figure 2. Similar inflammatory profiles in the brains and spinal cords of aged Optn^470T^ and WT mice, continued.** Representative RayBio® C-Series Mouse Inflammation Antibody Array C1 membranes incubated with brain lysates (left) and spinal cord lysates (right) from two-year-old male WT and Optn^470T^ mice are shown **(A)**. Mean values for individual proteins obtained by densitometric analyses ± SEM are shown for the brain **(B)** and spinal cord **(C)**. Three independent experiments were performed from the following groups of mice: 1) 4 WT and 5 Optn^470T^; 2)1 WT, 5 Optn^470T^; 3) 4 WT, 3 Optn^470T^. The data were analysed by Student’s t-test: * p<0.05.

**Suppl. Figure 3. T cell subsets in aged Optn^470T^ were comparable to WT male mice, continued.** Absolute splenocyte numbers from one- and two-year-old male WT and Optn^470T^ mice are shown **(A)**. Population frequency (%) **(B)** and absolute numbers (No) **(C)** of B cells, and CD4^+^/CD8^+^ ratio **(D)** are shown. Spleens of one- and two-year-old WT and Optn^470T^ male mice were stained for T cell activation markers, and gated as CD44^+^CD38^+^, CD44^+^CD69^+^, and CD44^+^KLRG1^+^ populations. CD8^+^ **(E-F)** and CD4^+^ **(G-H)** frequencies and numbers are shown. Staining for the indicated cytokines upon PMA/ionomycin stimulation is shown as frequency and MFI in CD8^+^ cells **(I-J)** and CD4^+^ cells **(K-L)**. The data from 6-7 one-year- and two-year-old WT and Optn^470T^ mice are shown as means ± SEM and analysed by two-way ANOVA: * p<0.05, ** p<0.01, ***p<0.001.

**Suppl. Figure 4. T cell subsets in aged Optn^470T^ were comparable to WT female mice.** Absolute splenocyte numbers from one- and two-year-old female WT and Optn^470T^ mice are shown **(A)**. Population frequency (%) and absolute number (No) of B cells **(B-C)**, T cells **(E** and **I)**, CD8^+^ and CD4^+^ **(F** and **J),** and CD4^+^/CD8^+^ ratio **(D)** are shown. Frequencies and numbers for CD8^+^ naïve (T_N_), central memory (T_CM_), and effector memory (T_EM_) **(G** and **K)**, and CD4^+^ T_N_, T_CM,_ and T_EM_ **(H** and **L)**. Spleens of one- and two-year-old WT and Optn^470T^ females were stained for T cell activation markers, and gated as CD44^+^CD38^+^, CD44^+^CD69^+^, and CD44^+^KLRG1^+^ populations. CD8^+^ **(M-N)** and CD4^+^ **(O-P)** frequencies and numbers are shown. Staining for the indicated cytokines upon PMA/ionomycin stimulation is shown as frequencies and MFI for CD8^+^ **(Q-R)** and CD4^+^ **(S-T)** cells. The graph shows the MFI of FOXP3 in Optn^470T^ normalized to one- and two-year-old WT female mice in CD4^+^FOXP3^+^CD25^+^ Tregs **(U)**. The graphs show CD25^+^ and CD25^-^ Tregs as frequencies **(V)** and numbers **(W)**. The data from 4-8 one-year- and two-year-old WT and Optn^470T^ mice are shown as means ± SEM and analysed by two-way ANOVA: * p<0.05, ** p<0.01, ***p<0.001, ****p<0.0001.

**Suppl. Figure 5. Innate immune characterization of WT and Optn^470T^ female mice.** Spleens of one- and two-year-old WT and Optn^470T^ female mice were stained for innate immune markers. Population frequency (%) and absolute cell numbers (No) are shown as following: frequency of non-T/non-B cells **(A)**, conventional dendritic cells (cDc) and macrophages (MΦ) **(B)**, and neutrophils and NK cells **(C)**; numbers for non-T/non-B **(D),** cDc and MΦ **(E)**, and neutrophils and NK cells **(F)**. The MFI for indicated activation markers normalized to one- and two-year-old WT and Optn^470T^ mice are shown for cDc **(G)**, and MΦ **(H)**. The data from 4-8 one-year- and two-year-old WT and Optn^470T^ mice are shown as means ± SEM and analysed by two-way ANOVA: * p<0.05, ** p<0.01

**Suppl. Figure 6. Immune cells isolated from the aged brains showed high autofluorescence.** Gating strategy for the immune cells isolated from the brains of two-month and one-year-old mice left unstained or stained for the indicated markers analysed by BD FACSAria^TM^ III cytometer **(A)**. Overlaid histograms show MFI from unstained and stained two-month-old, and unstained one-year-old immune cells in red (left) and green (right) channels **(B)**. Unstained immune cells isolated from two-month-old (left) and one-year-old (right) brains were seeded on coverslips, fixed, stained with DAPI, and analysed on Olympus IX83 fluorescent microscope **(C)**.

## Notes

### Competing Interest Statement

The authors have declared no competing interest.

