## Supplemental figures for "Neuroimmune characterization of optineurin insufficiency mouse model during ageing"

#### Supplemental material

### Neuroimmune characterization of optineurin insufficiency mouse model during ageing

Nikolina Mohovic[1]*, Josip Peradinovic[1]*, Andrea Markovinovic[1,2], Raffaello Cimbro[1], Zeljka Minic[3], Marin Dominovic[3], Hrvoje Jakovac[4], Jerneja Nimac[5,6], Boris Rogelj[5,7], and Ivana Munitic[1]

[1]Laboratory for Molecular Immunology, Department of Biotechnology, University of Rijeka, Radmile Matejcic 2, 51000 Rijeka, Croatia.

[2]Department of Basic and Clinical Neuroscience, Maurice Wohl Clinical Neuroscience Institute, Institute of Psychiatry, Psychology, and Neuroscience, King's College London, 5 Cutcombe Road, SE5 9RX, London, UK.

[3]Department of Biotechnology, University of Rijeka, Radmile Matejcic 2, 51000 Rijeka, Croatia

[4]Department of Physiology and Immunology, Medical Faculty, University of Rijeka, Brace Branchetta 20, 51000 Rijeka, Croatia.

[5]Department of Biotechnology, Jozef Stefan Institute, SI-1000 Ljubljana, Slovenia.

[6]Graduate School of Biomedicine, Faculty of Medicine, University of Ljubljana, SI-1000 Ljubljana, Slovenia

[7]Faculty of Chemistry and Chemical Technology, University of Ljubljana, SI-1000 Ljubljana, Slovenia

*These authors contributed equally.

#### Supplemental Figure 1.

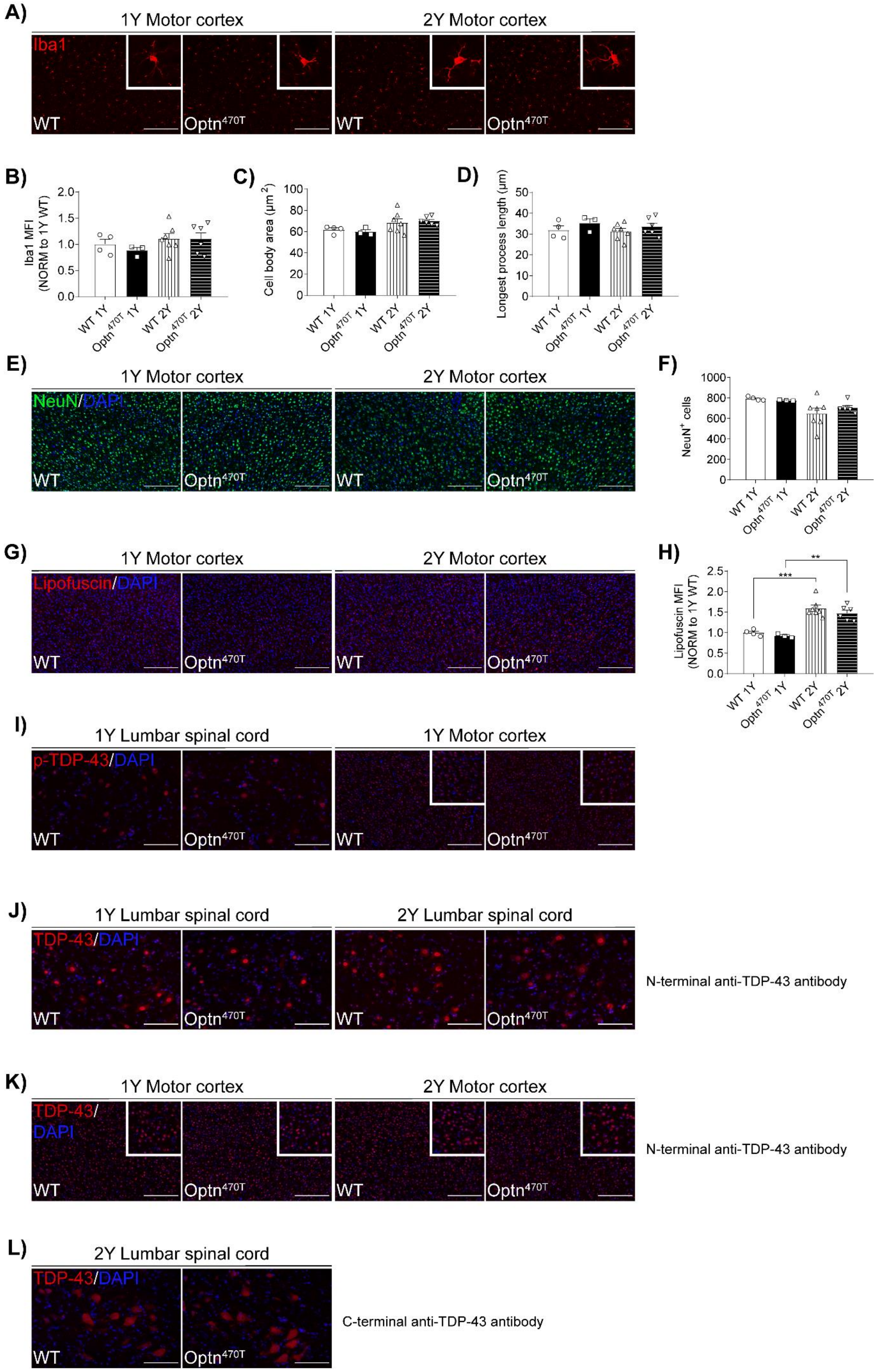

#### Supplemental Figure 2.

A)

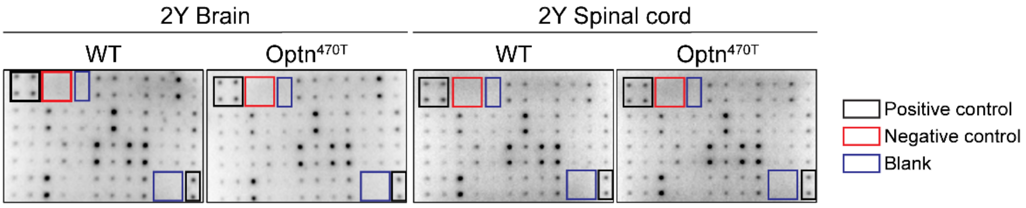


#### B) Brain

| | CXCL13 | CD30 | CCL11 | CCL24 | FasL | Fractalkine | G-CSF |
|---|---|---|---|---|---|---|---|
| WT | 1.2 ± 0.14 | 1.08 ± 0.16 | 0.6 ± 0.08 | 1.33 ± 0.20 | 0.93 ± 0.16 | 3.02 ± 0.28 | 0.28 ± 0.04 |
| Optn$^{470T}$ | 1.18 ± 0.17 | 0.55 ± 0.19 | 0.41 ± 0.17 | 0.92 ± 0.12 | 0.66 ± 0.16 | 2.54 ± 0.20 | 0.28 ± 0.04 |

| | GM-CSF | IFN-γ | IL-1α | IL-1β | IL-2 | IL-3 | IL-4 | IL-6 | IL-9 | IL-10 | IL-12 | IL-12p70 |
|---|---|---|---|---|---|---|---|---|---|---|---|---|
| WT | 0.49 ± 0.02 | 0.51 ± 0.09 | 1.36 ± 0.15 | 0.08 ± 0.05 | 0.29 ± 0.05 | 0.3 ± 0.06 | 2.96 ± 0.33 | 0.34 ± 0.04 | 1.12 ± 0.13 | 0.29 ± 0.04 | 0.49 ± 0.05 | 0.96 ± 0.1 |
| Optn$^{470T}$ | 0.46 ± 0.07 | 0.44 ± 0.08 | 1.44 ± 0.15 | 0.18 ± 0.01 | 0.44 ± 0.03 | 0.37 ± 0.05 | 2.72 ± 0.37 | 0.35 ± 0.08 | 0.91 ± 0.13 | 0.17 ± 0.07 | 0.51 ± 0.11 | 0.96 ± 0.09 |

| | IL-13 | IL-17A | I-TAC | CXCL1 | Leptin | LIX | XCL1 | CCL2 | M-CSF | CXCL9 | CCL3 | MIP-1γ |
|---|---|---|---|---|---|---|---|---|---|---|---|---|
| WT | 0.32 ± 0.005 | 0.29 ± 0.05 | 0.08 ± 0.04 | 0.56 ± 0.06 | 0.14 ± 0.01 | 2.54 ± 0.32 | 0.87 ± 0.05 | 2.6 ± 0.24 | 2.81 ± 0.23 | 0 07 ± 0 04 | 0.33 ± 0.08 | 0.26 ± 0.12 |
| Optn$^{470T}$ | 0.45 ± 0.05 | 0.39 ± 0.07 | **0.34 ± 0.05*** | 0.74 ± 0.04 | 0.26 ± 0.04 | 2.58 ± 0.11 | 0.96 ± 0.09 | 2.44 ± 0.29 | 2.54 ± 0.38 | 0.17 ± 0.08 | 0.51 ± 0.05 | 0.37 ± 0.04 |

| | CCL5 | CXCL12 | TCA-3 | CCL25 | TIMP-1 | TIMP-2 | TNF-α | TNF RI | TNF RII |
|---|---|---|---|---|---|---|---|---|---|
| WT | 0.3 ± 0.02 | 0.66 ± 0.13 | 2.13 ± 0.43 | / | / | 0.48 ± 0.09 | 0.59 ± 0.07 | 1.02 ± 0.12 | 0.44 ± 0.18 |
| Optn$^{470T}$ | 0.47 ± 0.16 | 0.9 ± 0.1 | 2.62 ± 0.19 | / | / | 0.65 ± 0.05 | 0.82 ± 0.09 | 1.23 ± 0.09 | 0.62 ± 0.14 |

#### C) Spinal cord

| | CXCL13 | CD30 | CCL11 | CCL24 | FasL | Fractalkine | G-CSF |
|---|---|---|---|---|---|---|---|
| WT | 1.5 ± 0.26 | 1.37 ± 0.22 | / | 1.38 ± 0.29 | 0.92 ± 0.27 | 1.86 ± 0.66 | 0.34 ± 0.18 |
| Optn$^{470T}$ | 1.81 ± 0.32 | 1.18 ± 0.11 | / | 1.37 ± 0.15 | 0.69 ± 0.12 | 1.59 ± 0.19 | 0.33 ± 0.07 |

| | GM-CSF | IFN-γ | IL-1α | IL-1β | IL-2 | IL-3 | IL-4 | IL-6 | IL-9 | IL-10 | IL-12 | IL-12p70 |
|---|---|---|---|---|---|---|---|---|---|---|---|---|
| WT | 0.58 ± 0.06 | 0.63 ± 0.07 | 1.5 ± 0.15 | 0.86 ± 0.14 | 0.2 ± 0.10 | 0.21 ± 0.12 | 3.74 ± 0.56 | / | 0.84 ± 0.15 | / | 0.63 ± 0.11 | 1.07 ± 0.23 |
| Optn$^{470T}$ | 0.6 ± 0.01 | 0.86 ± 0.1 | 2.00 ± 0.49 | 1.19 ± 0.30 | 0.43 ± 0.25 | 0.43 ± 0.20 | 3.93 ± 0.53 | / | 1.02 ± 0.15 | / | 0.69 ± 0.07 | 0.96 ± 0.1 |

| | IL-13 | IL-17A | I-TAC | CXCL1 | Leptin | LIX | XCL1 | CCL2 | M-CSF | CXCL9 | CCL3 | MIP-1γ |
|---|---|---|---|---|---|---|---|---|---|---|---|---|
| WT | 0.6 ± 0.06 | 0.5 ± 0.09 | 1.05 ± 0.16 | 0.89 ± 0.07 | / | 3.6 ± 0.53 | 0.98 ± 0.05 | 3.22 ± 0.38 | 3.6 ± 0.49 | / | 0.64 ± 0.06 | 0.54 ± 0.11 |
| Optn$^{470T}$ | 0.69 ± 0.01 | 0.68 ± 0.17 | 1.32 ± 0.28 | 1.18 ± 0.29 | / | 3.96 ± 0.42 | **1.32 ± 0.1*** | 3.33 ± 0.28 | 3.45 ± 0.40 | / | 0.71 ± 0.01 | 0.32 ± 0.03 |

| | CCL5 | CXCL12 | TCA-3 | CCL25 | TIMP-1 | TIMP-2 | TNF-α | TNF RI | TNF RII |
|---|---|---|---|---|---|---|---|---|---|
| WT | 0.91 ± 0.02 | 1.43 ± 0.25 | 4.1 ± 0.61 | / | / | 0.99 ± 0.24 | 1.27 ± 0.21 | 1.18 ± 0.19 | 2.14 ± 0.36 |
| Optn$^{470T}$ | 1 ± 0.04 | 1.66 ± 0.24 | 4.35 ± 0.54 | / | / | 0.83 ± 0.07 | 1.02 ± 0.1 | 0.69 ± 0.1 | 1.39 ± 0.21 |

#### Supplemental Figure 3.

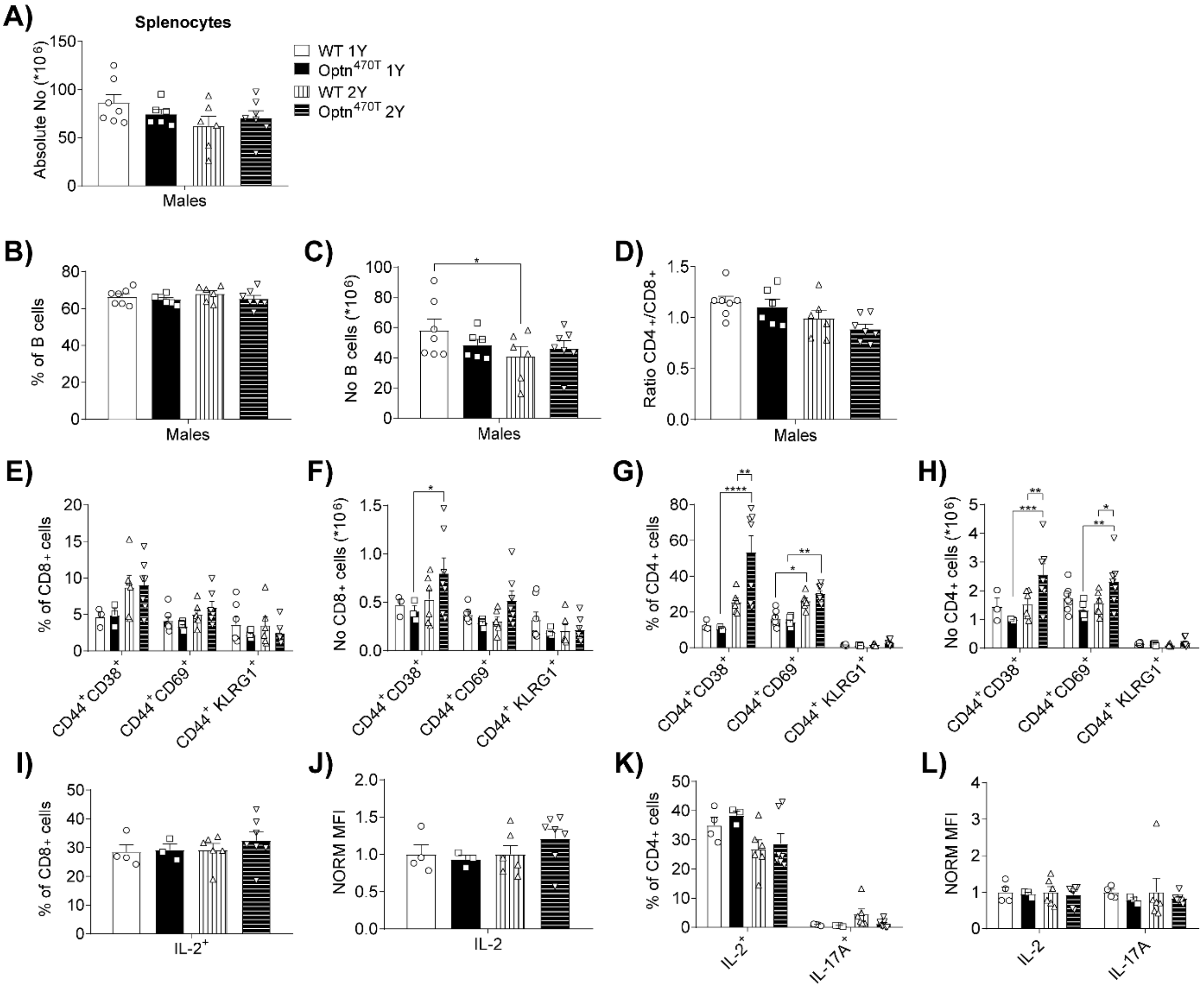

#### Supplemental Figure 4.

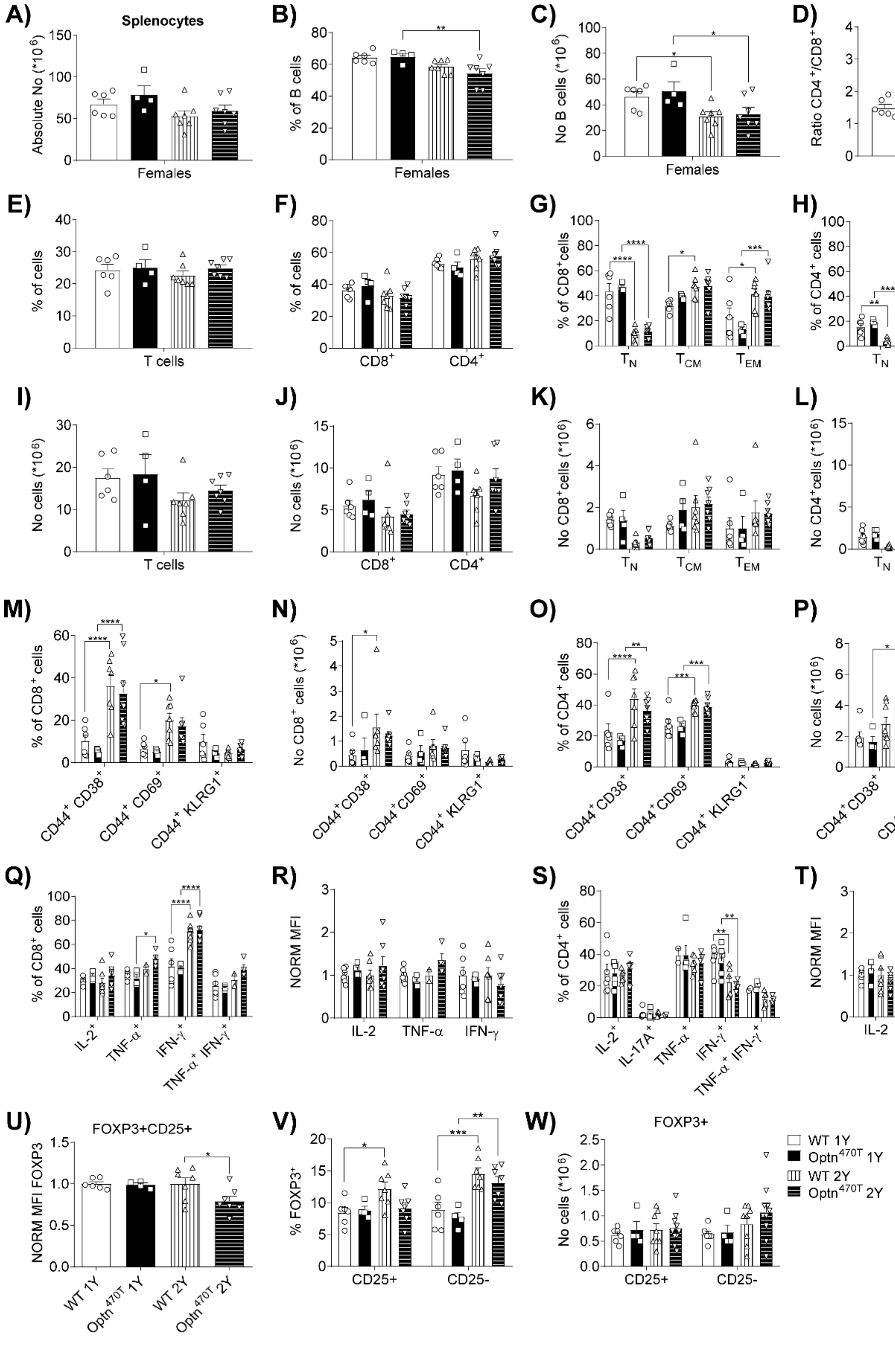

**Supplemental Figure 5.**

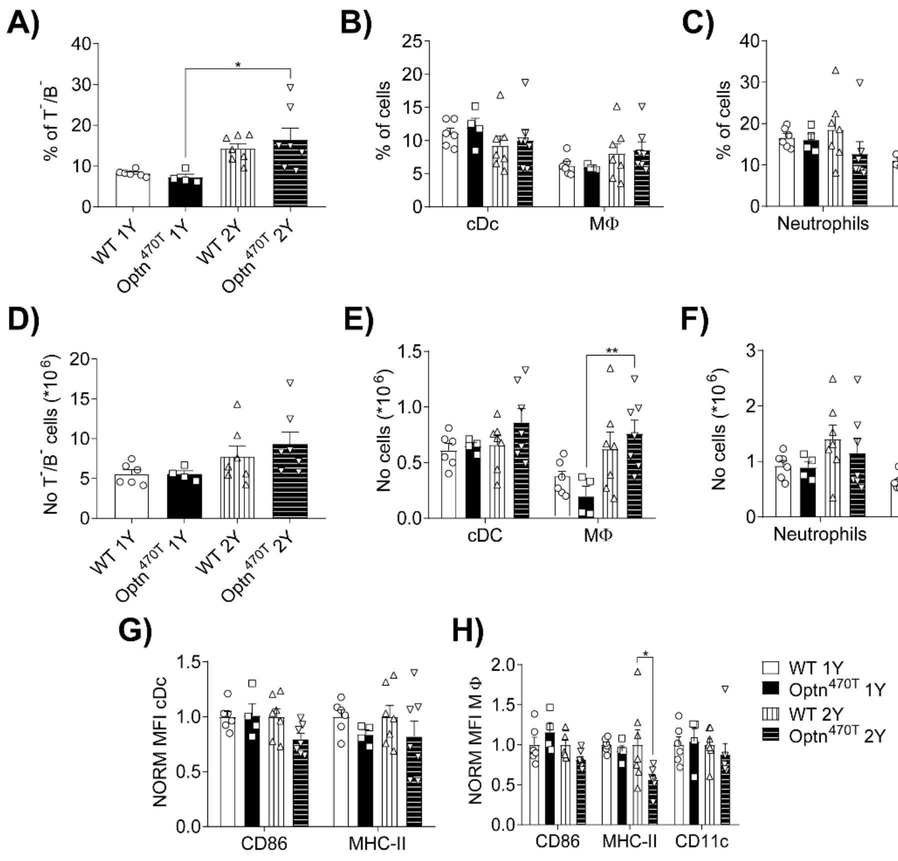

**Supplemental Figure 6.**

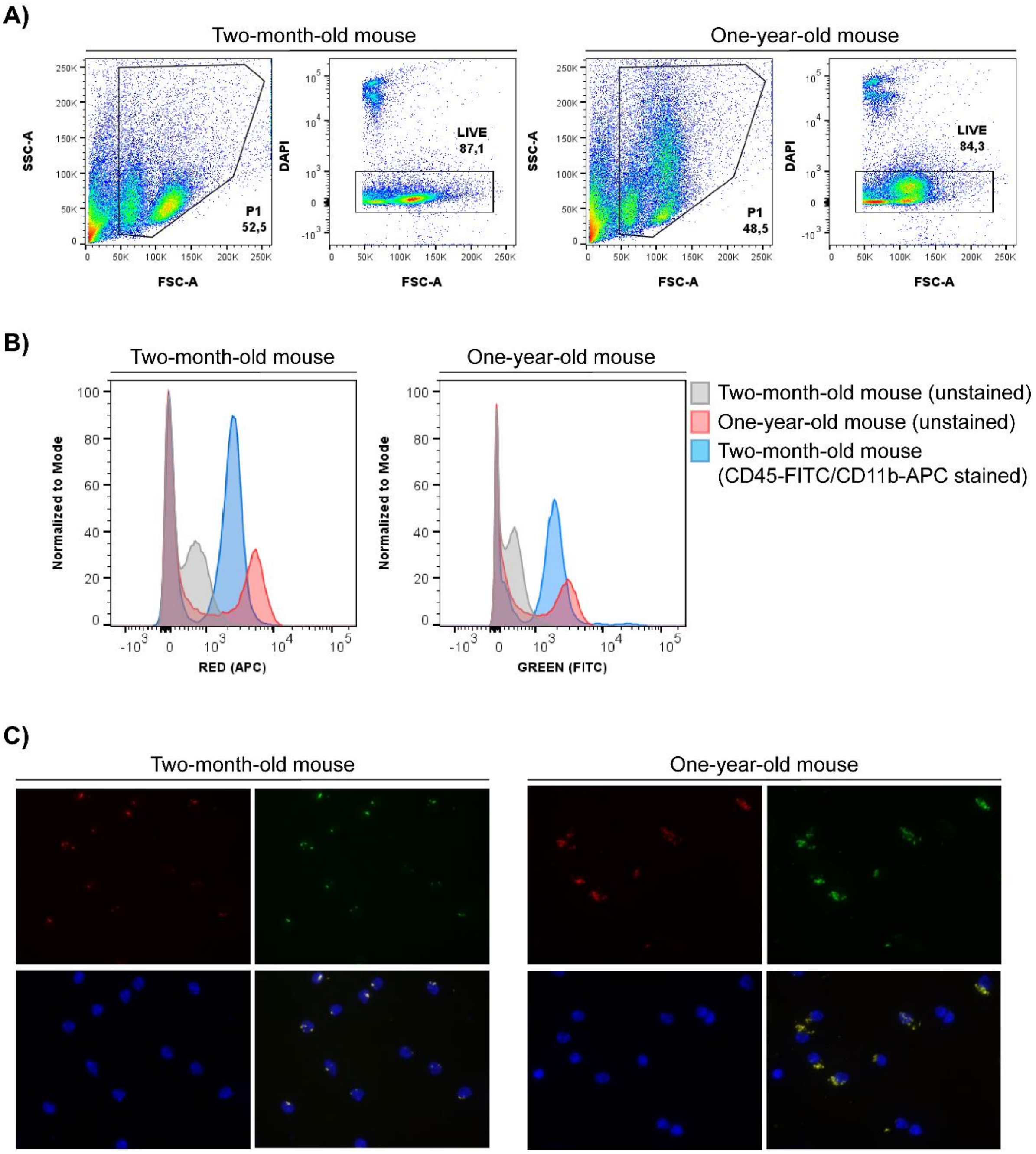
